# Enhancing Volatile Fatty Acid Accumulation in Seaweed-Arrested Anaerobic Digestion via a Two-Tier Framework of Engineering Diagnostics and Microbial Surveillance

**DOI:** 10.1101/2025.10.06.680780

**Authors:** Zhiyuan Zhao, Siman Liu, Hasan Nikkhah, Yidan Zhang, Yi Wang, Jiayi Liang, Shan Lu, Burcu Beykal, Mingyu Qiao, Baikun Li

## Abstract

This study presents a two-tier framework for brown seaweed–arrested anaerobic digestion (SW-AAD) by coupling engineering diagnostics (tier-1) with microbial surveillance (tier-2) to transform conventional digesters into volatile fatty acid (VFAs) biorefineries. In 70-day lab-scale batch tests (volume: 2 L), engineering diagnostics revealed that sequential additions of brown seaweed elevated salinity to 3.3% ash content, stabilized pH at 5.8–6.1, collapsed the CH₄/CO₂ ratio from 2.98 to <0.04, suppressed biogas by 96%, and boosted carbon-conversion efficiency from 10% to 52%. More than half of the influent carbon was redirected from methane to a liquid-phase VFAs pool, dominated by butyrate, hexanoate, and acetate, peaking at 14.5 g L⁻¹. Microbial surveillance using 16S rRNA sequencing presented a 10-fold decline in methanogens, alongside an increase in salt- and acid-tolerant acidogenic microbial families, including *Lachnospiraceae*, *Ruminococcaceae*, and *Clostridiaceae,* as well as seaweed-derived microbial families, including *Psychromonadaceae*, and *Marinomonadaceae*. Temporal synchrony and asynchrony-resolved analysis confirmed a strong correlation between two tiers, revealing an operational window featured with two patterns: a rapid engineering parameter response (where changes in engineering parameters were followed by a decline in methanogens within 0–3 days) and a slower microbial restructuring (where the surge in VFA concentration lagged the enrichment of seaweed-derived acidogens by 1–5 days). Based on this operation window, feedback control strategies were simulated for SW-AAD, indicating that cumulative VFAs (cVFAs) yield could be bolstered from 11.38 to 16.13 g COD L⁻¹ (+41.8%). A techno-economic assessment (TEA) revealed that this VFA-targeted SW-AAD can reduce capital expenditure by ∼20% compared to conventional AD. This study underscores the promise of asynchrony-resolved analysis of SW-AAD systems for robust and economically viable VFAs production from organic waste.

**Graphics:** 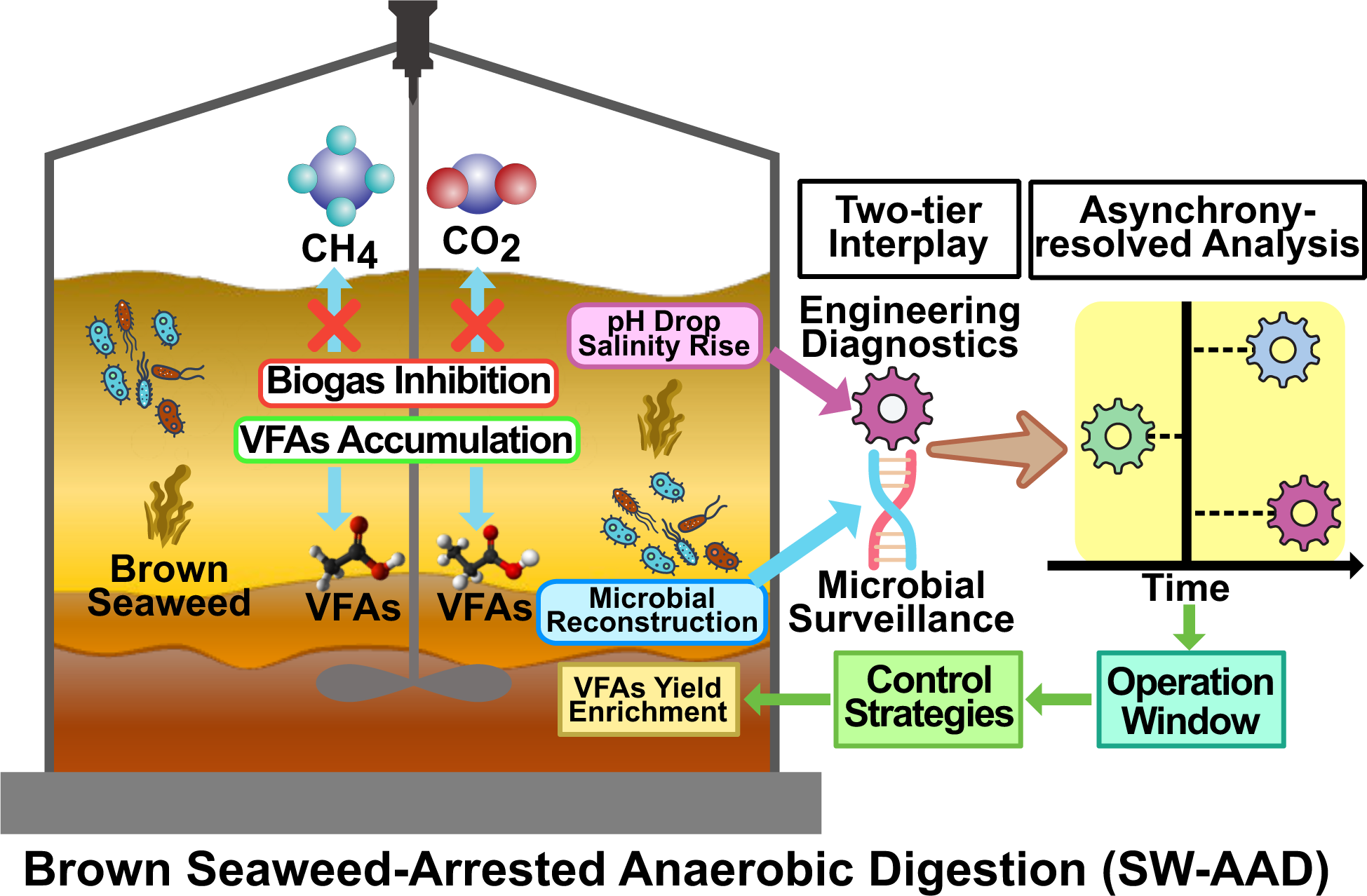

**Environmental synopsis:** Brown seaweed–arrested anaerobic digestion (SW-AAD) leverages seaweed-driven salinity, and acidification to suppress methanogenesis, diverting >50% of carbon to VFAs (14.5 g COD L⁻¹) while cutting methane by 99%. A two-tier framework couples engineering diagnostics with microbial surveillance and asynchrony-resolved analysis, enabling feedback control and low-emission waste valorization in digesters.

## Introduction

Anaerobic digestion (AD) in wastewater resource recovery facilities (WRRFs) converts organic waste (e.g., waste sludge, food waste)^1^ to biogas, primarily composed of methane (CH₄) and carbon dioxide (CO₂), providing a renewable source of clean energy^2^. Despite its benefits, AD faces limitations, including susceptibility to feedstock fluctuation^3^, long retention time (over 90 days)^3^, and poor energy utilization efficiency (2.1–12.7%)^4^. Arrested anaerobic digestion (AAD), a modified form of AD, redirects carbon to volatile fatty acids (VFAs, e.g., acetic, propionic, butyric and valeric acids) as the precursors for high-valued biofuels and biochemicals instead of methane,^5,6^ cutting retention time to 10–20 days and operating costs by 30%^7,8^. However, scaleup of AAD has been constrained by reliance on synthetic inhibitors (e.g., chloroform) that raise costs^9^, introduce contaminants to the final effluents^10^, and intensify petrochemical-derived carbon footprint^11^. Biomass-based inhibitors can address these problems by enabling AAD co-digestion. Especially, seaweeds (macroalgae) supply salinity (≤3.5% ash content) and lower pH (∼4.5) that curb methanogens and steer carbon to VFAs^11–13^. Previous studies showed that red seaweeds (e.g., *Asparagopsis taxiformis*) reduce ruminant enteric methane by 97%.^14^ However, slow growth rate (1.88%-3.30% per day)^15^, ecological requirement (e.g., warm tropical waters, symbiotic bacteria)^16^, and high production cost ($1 million per wet tonne) limit red seaweeds’ scalability^17^. In contrast, brown seaweeds (e.g., sugar kelp), which account for 34.65% of global cultivation ^18^, exhibit rapid growth rates (∼8.5% per day), low production costs ($200–300 per wet tonne) ^19^, and serve as non-food-grade biomass for fuel and fertilizer applications ^2022^. Brown seaweed AAD (SW-AAD) possesses a potential for sustainable VFAs production in a cost-effective manner^21^.

The mechanisms underlying SW-AAD systems are multifaceted. Acidification and elevated salinity impair methanogenic activity, damage cellular structures, and suppress CH₄ formation^22^. From an engineering perspective, engineering parameter (e.g., biogas production rate, pH, VFAs, and salinity) have been used to track the transition from AD to AAD^23^ ^24^. From a microbial perspective, seaweed-based chemicals directly inhibit methyl-coenzyme M reductase, a key enzyme in the Wolfe cycle of methanogenesis^25–30^. Yet microbiome restructuring during the transition from AD to AAD remains poorly resolved, limiting our ability to maximize VFAs yield. Moreover, single-perspective analyses (either engineering or microbial) are insufficient as they cannot capture the system’s inherent asynchrony (Figure 1 a-c): seaweed addition instantaneously alters engineering parameters like pH and salinity, while microbial community reconstruction is a gradual process with discernible changes emerging after days, creating an obvious asynchrony response system (Figure 1 c). This time-scale discrepancy reveals the importance of quantitatively capturing the dynamic and interplay between engineering parameters and the microbial community for optimization of SW-AAD systems.

**Figure. 1.**
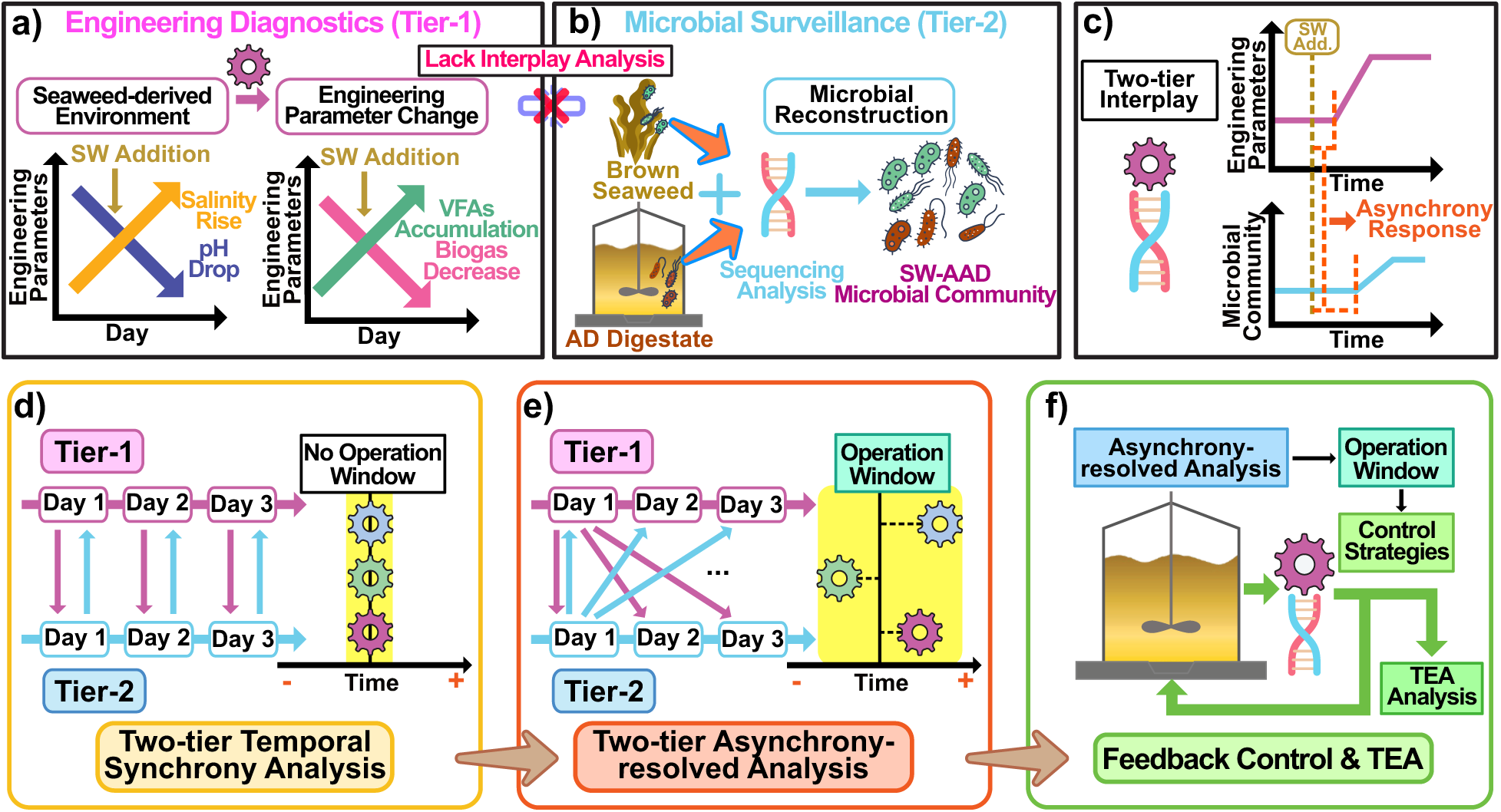
Two-tier, engineering diagnostics-microbial surveillance framework for SW-AAD system: from two-tier temporal synchrony and asynchrony-resolved analysis to feedback control and TEA; a) Engineering Diagnostics (Tier-1) - Periodic brown seaweed addition introduces seaweed-derived environment and change engineering parameters (pH, salinity), suppressing biogas production and enhancing VFAs accumulation, without interplay analysis, parameters are not yet linked with microbial; b) Microbial Surveillance (Tier-2) - Microbial surveillance captures the microbial reconstruction by sequencing analysis—methanogen suppression, acidogen enrichment, and reassembly into an SW-AAD community, without temporal analysis, linkage to engineering signals remains unresolved; c) Two-Tier Interplay. Illustration of the asynchronous response between engineering parameters and microbial community. (d) Two-Tier Temporal Synchrony Analysis: Direct and sequential correlations between the data obtained from two tiers on the same day (e.g., Tier 1 Day 1 vs. Tier 2 Day 1), and no operation window; e) Two-Tier Asynchrony-Resolved Analysis. Asynchrony: Cross and non-sequential correlation between the data obtained from two tiers on different days (e.g., Tier-1 Day 1 vs. Tier-2 Day 2), and an operational window established. f) Applying feedback control strategies during operational window for SW-AAD system optimization.

Recognizing these critical gaps of SW-AAD, we built a two-tier framework that fuses engineering diagnostics (Tier-1) with microbial surveillance (Tier-2) to quantify the interplay (Figure 1 a-b), identify the key triggers of methanogenic inhibition, and quantify the system’s asynchrony (Figure 1c). Specifically, the brown seaweed–induced salinity and acidification were captured by engineering diagnostics through engineering parameters (e.g., biogas production rate, pH, oxidation-reduction potential (ORP), ash content, conductivity and nutrient (e.g., C/N ratio)) that can reflect the transition from methanogenesis to acidogenesis (Figure 1a). In parallel, seaweed-derived community restructuring under acidification and osmotic stress after brown seaweed addition was captured by microbial surveillance that triggered methanogen suppression and acidogen enrichment (Figure 1b). To analyze the two-tier interplay, we first used temporal synchrony analysis to establish a statistical correlation between the tiers (Figure 1d), followed by asynchrony-resolved analysis to quantify the asynchrony and define the operational window (Figure 1e). Subsequently, we simulated the feedback control strategy based on our experimental data to adjust key operational variables (e.g., hydraulic retention time (HRT), internal circulation, pH range, and VFAs withdrawal) during the operation window (Figure 1f). This control strategy was expected to ensure stable and predictable performance of SW-AAD systems, providing a foundation for Techno-Economic Analysis (TEA), which remained limited for SW-AAD systems^31^ (Figure 1f).

Our objective was to develop a two-tier framework capable of quantifying the dynamic correlations between engineering parameters and microbial communities, characterizing the asynchrony responses to seaweed addition, and translating this information into operation window for feedback-control strategies with the ultimate goal of improving VFA yields in SW-AAD systems. Specifically, we systematically analyzed a series of engineering parameters and characterized microbial community during the transition period (70 days) of AD to AAD after brown seed additions. Based on the experimental data, we first applied temporal synchrony analysis to confirm the correlations and then deployed the asynchrony-resolved analysis to quantify the asynchrony extent and define the operational window. Subsequently, we simulated feedback control strategies within this operation window and determined the improvement of VFA production in SW-AAD using feedback control. Finally, we performed TEA to compare the SW-AAD with conventional AD in terms of feedstock source, operational costs, and product economic viability. This study turned the two-tier framework into operationally practical actions that improve the robustness and resilience of SW-AAD while maintaining high VFA yields.

## Methods and Materials

### Lab-Scale SW-AAD Bioreactor Experiment and Characterization

A lab-scale SW-AAD experiment was conducted in a BIOFLO 110 Fermentor (New Brunswick Scientific, NJ) maintained at 35 °C. Digestate was sourced from a 5,600 m³ co-digestion reactor (25-day HRT) at the Greater Lawrence Sanitary District (GLSD) Wastewater Treatment Plant (North Andover, MA; 190,000 m³/day capacity). Sugar kelp (*Saccharina latissima*) was cultivated and harvested in April 2023 from Fishers Island Sound (CT, USA), processed into slurry, and stored at −20 °C for ∼1 year. The Fermentor (11 cm diameter × 25 cm height, 2 L volume; Figure S1) was agitated at 120 rpm using a 6-blade impeller (2 cm diameter, 1.75 cm width). All fittings were sealed with DuPont MOLYKOTE High-Vacuum Grease (Fisher Scientific, NH) to ensure strict anaerobic. After sealing, the reactor was purged with 99.95% nitrogen gas (Airgas Co., CT). Biogas was vented through a headspace outlet connected to a fume hood (Labconco, MO).

The SW-AAD bioreactor was operated as a fed-batch system for 70 days, during which the transition from conventional AD to a VFA-accumulating state was observed. As a fed-batch experiment, the system did not have a defined HRT. By contrast, in the control simulation and in full-scale continuous AD/AAD systems, HRT is essential because it sets microbial residence time and contact with acidic/saline conditions, thereby sustaining methanogen inhibition, preventing washout/over-acidification, and maximizing VFA productivity^32^.

Brown seaweed slurry (∼1 kg) was added on Days 8, 15, and 22, replacing a portion of the existing digestate. Throughout the experiment, the reactor was continuously stirred to ensure proper mixing until a quasi-steady state was achieved, which was defined by stable trends in pH, VFAs, and gas production over several days. To monitor this transition, 15 mL digestate samples were collected three times per week for a suite of chemical analyses, including pH, ORP, conductivity, VFAs, COD, NH₄⁺, salinity^33^ (Figure S1, Figure S2). Salinity was further analyzed via ash content measurements^34^ (Test S1), while the concentration and composition of VFAs were confirmed using Gas Chromatography-Mass Spectrometry (GC-MS)^35,36^ (Test S2-S4).

### 16S rRNA Gene Sequencing Analysis of Seaweed and Mixed Sludge

The digestate samples from different time points over the 70-day operational period and the seaweed samples were collected and stored at −80°C for the following procedures. DNA was extracted using MagAttract PowerSoil Pro DNA kit (Qiangen, Inc). DNA quantification was performed using the Quant-iT PicoGreen kit (Invitrogen, ThermoFisher Scientific). Partial bacterial 16S rRNA (V4) region was amplified using 515F (GTGYCAGCMGCCGCGGTAA) and 806R (GGACTACNVGGGTWTCTAAT) with Illumina adapters and dual indices^37^. The Polymerase Chain Reaction was incubated at 95°C for 2 minutes, then 30 cycles of 30s at 95.0°C, 30s at 50.0°C and 60s at 72°C, followed by final extension at 72°C for 10 minutes. The cleaned DNA was sequenced on the MiSeq using v2 2×250 bas pair kit (Illumina, Inc.).

Sequences were processed in Mothur 1.36.1, following the MiSeq standard operating procedure (SOP) as described by Kozich et al.^38^. The sequences were aligned against the SILVA non-redundant database, version 138.1. After demultiplexing and quality checking steps, the sequences were clustered into operational taxonomic units (OTUs) at 97% similarity. Alpha diversity, which reflects the richness and evenness of microbial communities within individual samples, and beta diversity, which evaluates differences in microbial community composition between samples^39^ were determined by subsampling to 10,000 reads per sample. Alpha and beta diversity were performed using vegan package in R (≥4.1.0)^40^. Microbial communities were examined across various taxonomic levels, and the top ten most relatively abundant taxa were identified through R-based computational screening. The relative abundance (*P_i_*) of each taxa is calculated in the formulation below^41^:

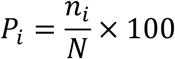

Where n_i_ is the number of individuals (counts or sequences) of the *i*-th taxon, N is the sum of all individuals across all taxa in the community. Methanogens were identified by isolating archaea taxa out and cross-referencing them with taxa commonly reported in other anaerobic digestion systems^42,43^. Similarly, bacteria taxa related to VFA production were identified based on previously reported VFA-producing organism in gut microbiota and anaerobic digestion environments^44^. Brown seaweed-derived bacteria were identified from a seaweed sample, and marine-derived taxa were validated by their presence in seaweed samples and absence in sludge samples.

### Establishing two-tier Interplay through Temporal Synchrony Analysis

We first performed temporal synchrony analysis between the two-tier (engineering diagnostics and microbial surveillance) to confirm that these two tiers are correlated (Figure 1d). Specifically, all data from these two tiers were aligned to the daily frequency (Day: 0–70) using shape-preserving monotone cubic interpolation (PCHIP) (Text S5) to avoid overshoot and preserve measured trends between samples (Table S4). Pearson cross correlations matrix was calculated at matched timestamps to quantify synchrony relationships, with results reported as Pearson *r* (Text S6).

### Quantifying Two-tier Interplay through Asynchrony-Resolved Analysis

Along with using temporal synchrony analysis to determine whether these two tiers have correlations, we also performed temporal asynchrony analysis to quantify the two-tier interplay. The temporal asynchrony is defined that the engineering parameters and microbial communities do not proceed simultaneously (Figure 1e). To quantify these asynchronous correlations and interplay, we focused on the periods following seaweed addition. The analysis involved a two-step quantification process. First, event-aligned cross-correlation (EACC) was used to explore the asynchrony relationship between an engineering parameter and a microbial community (Text S7)^45^. EACC could provide asynchrony magnitude. Second, bivariate Vector Autoregression (VAR)-Granger tests were used to verify and confirm the direction of these relationships (Text S7)^46^. VAR–Granger could provide more detailed directionality, which converts the two-tier interplay information into actionable operation windows for feedback control. The mechanisms of EACC and VAR-Granger tests are described in Table S3 and Figure S3.

### Executing Feedback Control Strategies in the Operation Window by a Data-driven Simulation

We subsequently translated the asynchrony-resolved analysis into an operation window for the feedback control strategies intended for continuous-flow digesters (e.g., CSTRs), where HRT is an important process variable governing biomass retention and stability^32^. Rapid variations in engineering parameters (e.g., acidic pH, salinity increase) were used to identify the arrested stage, while changes in the microbial community predicted subsequent changes in biochemical patterns, such as a decline in methanogens indicates a reduction of methane production or the proliferation of acidogens indicates an increase in VFA accumulation. Specifically, feedback control strategies were simulated in Python 3.11 based on our SW-AAD experimental data. Control strategies were triggered when engineering parameters or microbial communities passed the pre-defined thresholds obtained from our experimental data and/or the previously reported values. In this study, four primary control strategies were applied within the operational window, specifically (i) prolonging HRT by holding effluent for ∼24 h after salinity/pH shocks to sustain methanogen suppression^47^; (ii) applying an internal recirculation for one day to enhance mass transfer and substrate contact, thereby improving conversion of substrate and mitigating transport limitations^48^; (iii) maintaining a mildly acidic pH range (5.6-5.9) within the AAD system ^49^; and (iv) *in-situ* withdrawal 10% of VFAs when the system shows signs of over-acidification (pH<5.5) or product inhibition^50^. By withdrawing a small fraction of VFAs, we could mitigate inhibitory of VFA acidic products while preserving hydraulic and microbial stability. Details of the feedback control execution are explained in Figure S8, Text S8 and Table S7.

### Techno-Economic Analysis (TEA) of SW-AAD systems and AD systems

The TEA of the SW-AAD system with/without feedback control strategy versus the conventional AD system was conducted at the industrial scale, where all the systems were assumed to have a 20-year plant lifetime and an annual processing capacity of approximately 90,000 tons of feedstock (89,800 tons, the same capacity as the one we took digestate samples)^51^. The annualized cost of each system was calculated to ensure a consistent basis for comparison. Specifically, we assumed 330 operation days per year and an interest rate of 0.08, consistent with our previous study of TEA analysis ^52–54^. For the capital expenditure (CAPEX) and operational expenditure (OPEX) assessment of the AD biogas facility, we utilized the reported numbers from published studies^55^. Revenue streams were derived from electricity sales for AD, whereas VFAs for SW-AAD. Moreover, tipping fees (or gate fees) are an essential revenue stream for waste management facilities, often exceeding $100 per ton^56^, which is critical for covering operational costs such as personnel, maintenance, and regulatory compliance, as well as for financing recycling and infrastructure enhancements^57^. A tipping cost of $50 per ton was used for our analysis. Based on the balance of costs (CAPEX and OPEX) and revenues (electricity, VFAs, and tipping fees), the annual profit of each system was subsequently calculated. Details of the calculation methodology are provided in Text S9.

## Results and discussion

### Characterization of the SW-AAD System through Engineering Parameter Diagnostics

Bi-daily measurements of key engineering operational parameters over the 70-day operational period revealed a clear transition from conventional AD to SW-AAD following the sequential brown seaweed additions on Days 8, 15, and 22. At the initial phase (Days 0–7), the pH in the reactor was 7.78-7.83 (Figure 2a), indicative of operational stability typical for AD systems^58^. Following the transition phase, (first seaweed addition (Days 8–14) and second addition (Days 15-21)), the reactor exhibited moderate acidification, with pH descending slightly to 6.39 (Day 20), which was below normal pH range (6.5–7.8) for AD systems^59^. The pH subsequently stabilized between 5.60 – 6.00 after the third addition of seaweed (Days 22), turning the system into SW-AAD phase (after day 22). The drastic pH drop following brown seaweed addition implied an enhanced acidogenic activity, primarily driven by fermentation of seaweed slurry, which aligned with previous findings of pH declining from 7.5 to 7.2 in mesophilic digestion of *Ascophyllum nodosum* ^60^ and was likely caused by organic acids produced from fermentable carbohydrates (e.g., mannitol and laminarin).^61^ In the later operational period (Days 22-77), pH stabilized at ∼5.70-6.31, indicating that pH serves as a robust proxy for acidogenic performance.

**Figure. 2.**
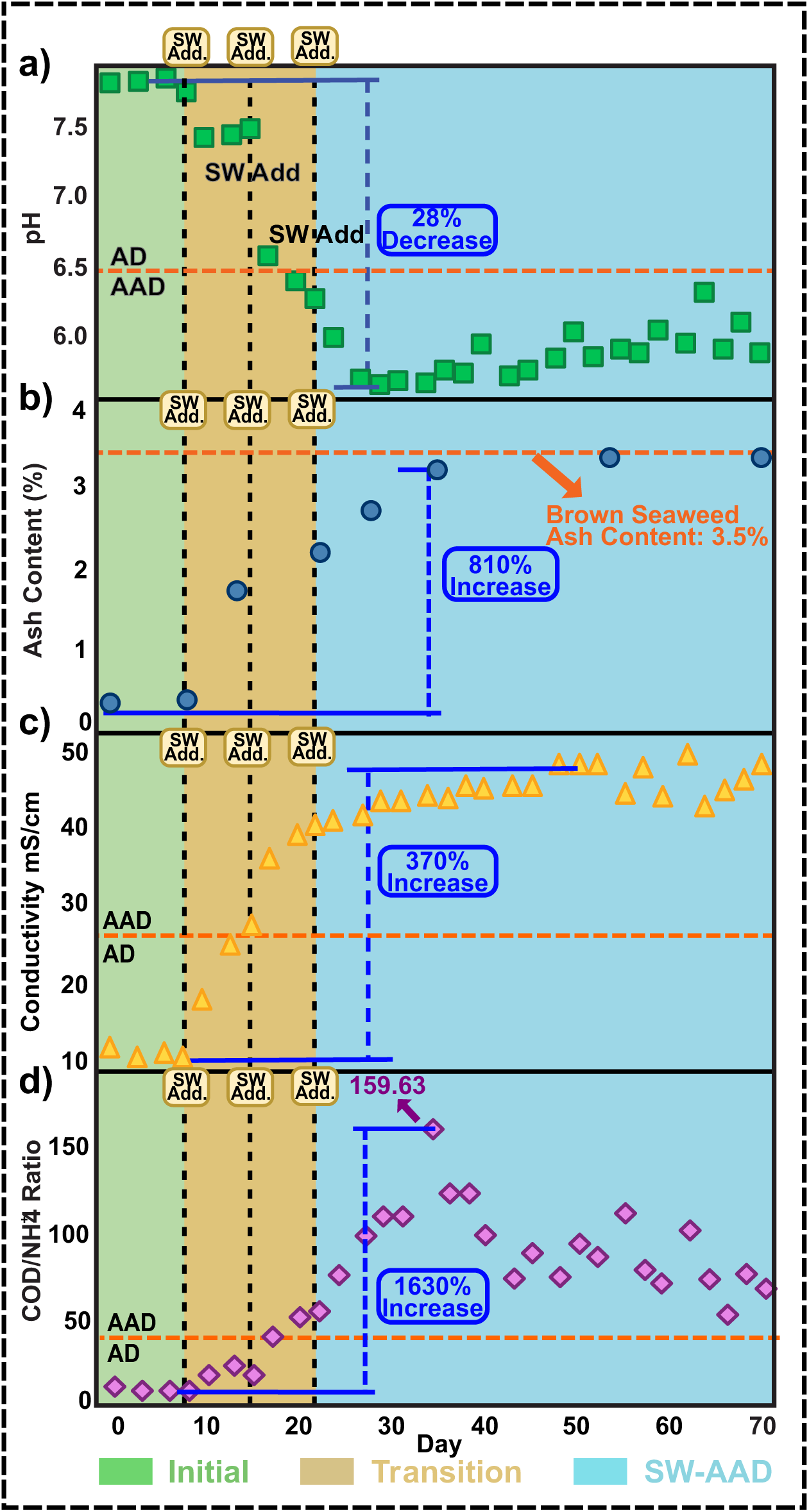
Engineering Diagnostics (Tier-1) capturing the transition from conventional AD to AAD across three phases: Initial (green), Transition (tan), and SW-AAD (blue). (a) pH dropped by 28% following sequential additions of brown seaweed, marking the onset of acidogenesis. (b) Ash content increased by 810% and stabilized at 3.5%, consistent with seaweed’s mineral load. (c) Conductivity rose by 370%, reflecting salt accumulation from seaweed biomass. (d) COD/NH₄⁺ ratio surged 1630% and peaked at 159.63 on Day 30, indicating strong carbon enrichment and nitrogen dilution. These parameters serve as engineering indicators of arrested methanogenesis and enhanced acidogenesis during the transition from AD to SW-AAD.

Maintaining a mildly acidic environment at pH value of 5.5–5.9 in SW-AAD can selectively suppress methanogenic archaea (optimal pH: 6.8–7.5) while allowing acidogenic bacteria to sustain metabolically active and redirecting carbon flux toward the accumulation of VFAs^62^. At mild acidification pH (5.5 – 6.0), methanogenic archaea are inhibited due to the disruption of key enzymes involved in hydrogenotrophic and acetoclastic methanogenesis.^63^ This metabolic imbalance prevents VFAs consumption and promotes their accumulation, contributing to the targeting intermediate product recovery in AAD systems. Furthermore, pH of 5.0–6.0 had been found to enrich acid-tolerant fermenters such as *Clostridium* spp. that efficiently channel carbon toward VFAs production, which further boost acetate, propionate, and butyrate yields.^64^ These findings demonstrate that in brown seaweed–aided operational conditions, maintaining a stable mildly acidic pH signifies an effective transition from AD to AAD.

In terms of salinity, the ash content (inorganic salts) increased from 0.4% to 3.3% following brown seaweed additions (Figure 2b), indicating a substantial rise in salt concentration. This ascending trend corresponded with an increase in conductivity from 10.8 mS/cm to 48.8 mS/cm in the reactor, confirming the elevation of ionic strength due to seaweed incorporation (Figure 2c). Previous studies had shown that elevated salinity disrupted osmotic balance, causing plasmolysis and energy stress that especially inhibit methanogens, while favoring halotolerant, acidogenic microbes.^65^ Furthermore, tremendous inhibition of methanogenesis had been found at NaCl concentrations as low as 8–13 g/L, with severe impairment or complete suppression observed beyond ∼20 g/L^66^. This sensitivity reflects inadequate osmoregulatory capacity of methanogens, leading to the vulnerability of enzymes and cellular machinery to osmotic stress^67^. By contrast, acidogenic bacteria endure high salinity by biosynthesizing or importing compatible solutes (e.g., small organic osmolytes) that stabilize proteins and maintain turgor and by remodeling membrane lipids to preserve fluidity under stress^68^. This selective pressure favors the enrichment of salt-tolerant taxa (e.g., *Psychromonadaceae*) which excrete broad polysaccharide-degrading enzymes, enabling sustained VFAs production and competitive advantage under saline, mildly acidic environments^69^. Along with pH and salinity, other engineering indicators also tracked the transition from AD to AAD. Specifically, the COD/NH₄⁺ ratios exceeded 30 (carbon-rich, N-limited), consistent with VFA build-up and instability^70^ (Figure 2d). COD in the AD reactor increased from 16,400 on Day 6 to 41,000 mg/L on Day 34 (Figure S1), while NH₄⁺ fell from 1,600 mg/L on Day 6 to 450 mg/L on Day 34 (Figure S4a), reflecting dilution by seaweed with negligible NH₄⁺^71^. Simultaneously, ORP ascended from ∼–240 to –130 mV (Figure S4c), approaching/exceeding the value (∼ –185 mV) associated with impaired methanogenesis ^72–74^ and reducing the activity of redox-sensitive enzymes (e.g., methyl-coenzyme M reductase) ^75^. By contrast, facultative fermenters and acid-tolerant anaerobes tolerate –250 to –100 mV^76,77^. Collectively, these changes act as clear, engineering-level markers for the seaweed-driven transition from AD to SW-AAD. Concurrently, biogas production fell from 1,080 mL d⁻¹ at ∼75% CH₄ (CH₄/CO₂ ratio: 2.98) to 96 mL d⁻¹ after the seaweed addition on Day 8, and to ≤0.72 mL d⁻¹ on Days 54–65 (−96% biogas; −99.98% CH₄; Figure 3a, S4d), consistent with the previous study findings (>90% inhibition at a seaweed polyphenol loading of 15%, based on volatile solids)^78^. The VFA was 1,037–1,282 mg COD L⁻¹ in the initial phase (Days 0–7) (Figure 3c), rose to 9,939 mg COD L⁻¹ after two additions of brown seaweed, peaked at 14,526 mg COD L⁻¹ on Day 40, and remained ∼10,377 mg COD L⁻¹ through Day 68. This upward trend corresponded with mild acidification (pH: 5.70–6.31) and ascending salinity (Figure 2a–b), indicating that osmotic/acid stress redirected carbon from methanogenesis towards VFA production in SW-AAD. Our system process also achieved a high VFA yield in comparison to previously reported values (Table S2).

**Figure 3.**
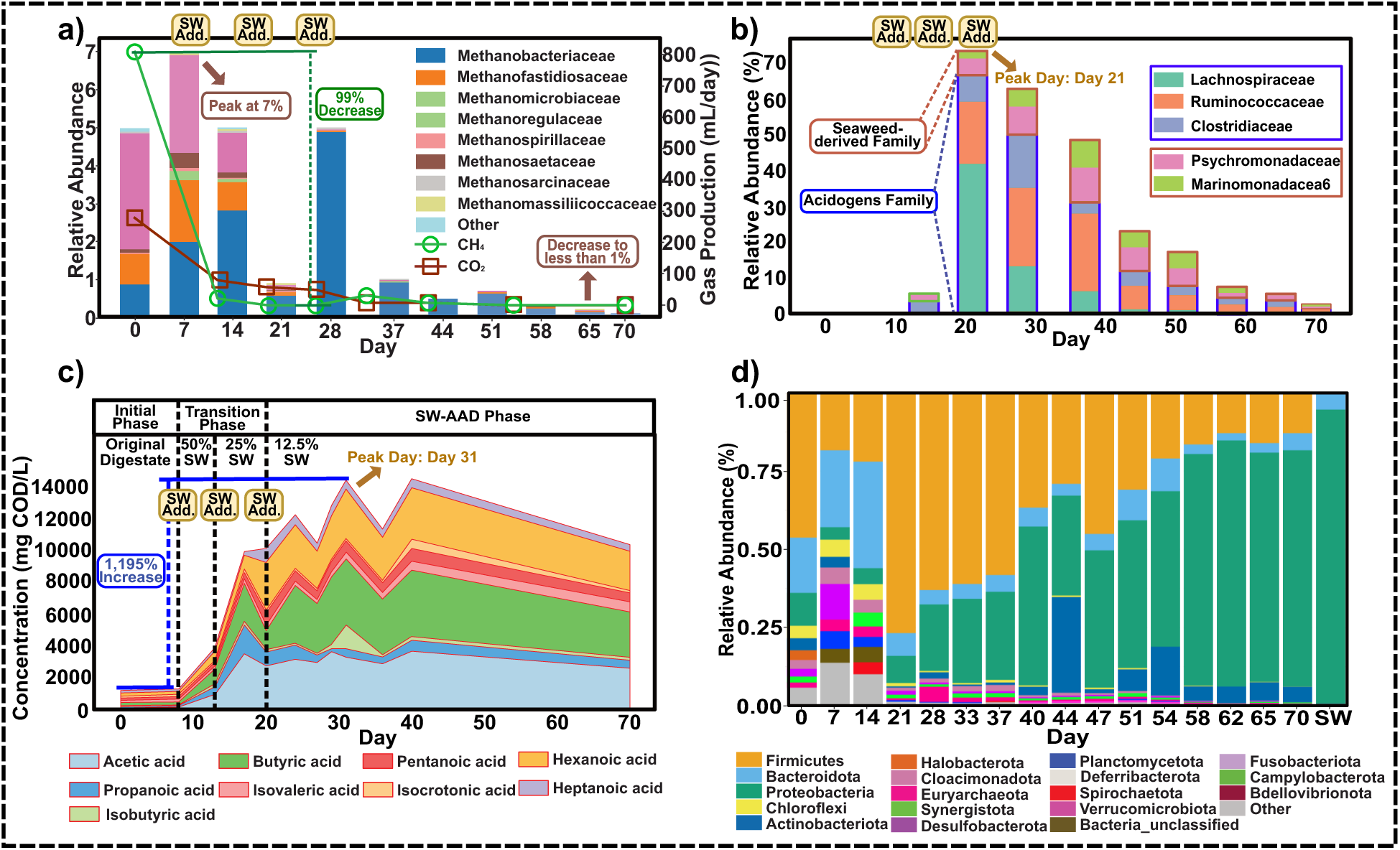
Microbial surveillance (Tier-2) along with biogas and VFAs yield to capture the transition from AD to SW-AAD. a) Relative abundance of methanogenic archaeal families (e.g., *Methanobacteriaceae*, *Methanomicrobiaceae*) and corresponding biogas production rates. Methanogen abundance peaked at ∼7% before seaweed addition, then declined by 99%, with CH₄ production falling below detectable levels. b) Relative abundance of key acidogenic microbial families, including *Lachnospiraceae*, *Ruminococcaceae*, *Clostridiaceae*; Seaweed-derived families, including *Psychromonadaceae*, and *Marinomonadaceae*. These families were substantially enriched after seaweed addition, peaking at over 70% relative abundance by Day 30. c) Temporal evolution of VFA composition across 70 days. Sequential additions of brown seaweed produced a 1,195% increase in total VFAs (1,000 to 14,000 mg COD L⁻¹), with butyric, hexanoic, and acetic acids dominating. d) Relative abundance of the top 10 bacterial and archaeal phyla in the AAD system from Day 0 to Day 70. Community shifted markedly after seaweed addition, with dominant phyla changing from *Euryarchaeota* and *Firmicutes* to *Bacteroidota* and *Proteobacteria* during the transition from AD to SW-AAD.

### Characterization of the transition of AD to SW-AAD through microbial dynamic surveillance

To elucidate the variation of the microbial community during the transition from AD to SW-AAD, we monitored the relative abundance of the microbial community at phylum levels (Figure 3d) and genus levels (Figure S7). Prior to seaweed addition (Day 0 ∼7), the microbial community was dominated by *Firmicutes* (31%∼46 %) and *Bacteroidota* (10%∼18 %), followed by *Proteobacteria* (11%∼43 %), which are fermentative microorganisms commonly found in anaerobic sludge^79^. Seaweed addition reshaped the microbial community in the AD system, as reflected in both alpha and beta diversity (Figure S6a and S6b). Alpha diversity measured by the inverse Simpson index(1/D)^80^, which declined from ∼ 40 to ∼15 after seaweed additions on Day 8, Day 15, and Day 22, indicating seaweed-induced environmental stress. With seaweed accounting for 87.5% of biomass in the system Day 22-51 (Figure S6a), the microbial community adapted, and diversity rebounded, enabling VFA-producing bacteria to proliferate. Beta diversity analysis revealed that seaweed addition altered the microbial community structure (Figure S6b), which progressively transitioned into a distinct community suited to the AAD system.

Seaweed addition introduced two marine-derived bacterial taxa--*Marinomonas* and *Psychromonas* (Figure S7). Specifically, *Marinomonas* was isolated from brown algae *Laminaria* and excrete alginate lyases that had been found to contribute to VFA production^81^. *Marinomonas* rose up to 28% of total community after three events of seaweed addition (Figure S7) and remained among the top ten genera until Day 58 (Figure S6a). On Day 23, the pH of the AAD system dropped below 6.0 (Figure 2a), which was beyond the optimal growth range (pH: 6.0∼8.0) for *Marimonas* ^82^ and resulted in the decline in its relative abundance^64^. Furthermore, *Psychromonas* species peaked at 11% on Day 33 (Figure S7) and were reported to hydrolyze proteins and lipids, leading to buildup of VFAs.^83^ Previous studies revealed that *Psychromonas* species secreted phospholipases, proteases, and peptidases, and could hydrolyze proteins and lipids extracellularly^84,85^. The introduction of these two seaweed-derived taxa had also been found to facilitate a VFA-producing microbial community in anaerobic sludge.^68,69^

All methanogens were confirmed to originate from the AD digestate, not the seaweed, as 16S rRNA sequencing detected no archaea species in the seaweed samples (Figure 3d). Following three seaweed additions on Day 8, 15 and 22, the seaweed proportion ascended to 87.5% of feedstock, while archaeal abundance (initially ∼5%) peaked at 7% in the transition phase (∼Day 7) and then declined to ≤1% by Day 33 (Figure 3a), despite methanogens comprising 98–100 % of archaea and being dominated by *Methanobacteriaceae, Methanofastidiosaceae*, and *Methanosaetaceae* (Figure 3a). The total ammonia nitrogen (TAN) remained less than 1.6 g/L (Figure S4a, Table S1), excluding the possibility that high ammonia inhibited methanogesis^86^ and implying that brown seaweed addition was the main reason for methanogen inhibition. Among these three methanogens, *Methanobacteriaceae* and *Methanosaetaceae* had been found to produce methane via acetoclastic pathway, whereas *Methanofastidiosaceae* employed a methylotrophic (methyl-CoM reduction) pathway^87^, meaning that the seaweed additions substantially reshaped the microbial community, selectively suppressed methanogen populations and promoting VFA accumulation.

The observed VFA accumulation was validated by monitoring VFA producing bacteria. Three bacterial families, *Lachnospiraceae, Ruminococcaceae, and Clostridiaceae,* were identified (Figure 3b), peaking at 66.75% around the third seaweed addition on Day 21 and then gradually decreasing to 1.11% on Day 70. Specifically, *Lachnospiraceae* had been known to produce VFAs^88^, and a genus under this family, *Epulopiscium*, was found in the SW-AAD system and became dominant (42.14% of the whole community) on Day 21 (Figure 3b). *Epulopiscium* species had been often found in marine tropical fish gut and contributed to VFA prodcution.^89^ On the other hand, *Ruminococcaceae*, associated with butyrate and propionate production ^82,90,91^, became dominant from Day 28, peaking at ∼21.80% (Figure 3a-3c).^64,73,74^ *Ruminococcaceae* is a family often found in human gut microbiota and was correlated to high propionate production in rumen environment.^92^ Furthermore, *Clostridiaceae*, previously identified as associated with butyric acid production in anaerobic fermentation ^93^, peaked at 14.80% on Day 28 (Figure 3b). Therefore, change in these bacterial families confirmed the microbial contribution to VFA accumulation in the SW-AAD system. From the observation of our experimental results, engineering parameters and microbial community are correlated with each other, but their changes are asynchrony. The peak relative abundance of acidogens was observed on Day 21(Figure 3b), preceding the maximum VFAs concentration on Day 31 by 10 days(Figure 3c). Therefore, to establish the interplay between the two tiers, we first use temporal synchrony analysis to confirm a statistical correlation, followed by asynchrony-resolved analysis to quantify the asynchrony.

### Temporal Synchrony Analysis for Establishing Two-tier Interplay in SW-AAD

Based on the experimental data of engineering parameters (Tier 1) and microbial communities (Tier 2), we performed temporal synchrony analysis to quantitatively confirm the statistical correlations and elucidate the two-tier interplay. The correlation matrixes confirmed the relationships driven by methanogen suppression, as measured by the Pearson correlation coefficient (ρ). A strong positive correlation was observed between pH and the dominant methanogens (ρ: 0.92–0.93), whereas COD, ash, and VFA levels were strongly anti-correlated (ρ: −0.87 to −0.95). Furthermore, CH₄ production was tightly coupled with the abundance of *Methanosaetaceae* (ρ: 0.97) and *Methanofastidiosaceae* (ρ: 0.86), while CO₂ exhibited an inverse relationship (Figure S5a-b). Conversely, salt- and acid-tolerant fermenting families, including *Ruminococcaceae*, *Clostridiaceae*, and seaweed-derived taxa (*Psychromonadaceae, Marinomonadaceae*), were positively associated with rising COD and VFA concentrations (ρ: 0.43–0.60) (Figure 4a). This correlation was also reflected in the VFAs composition, where butyrate, acetate, and hexanoate comprised 43% on day 1 and arose to ∼76% by Day 40. Specific microbial families were linked to the surge in VFAs, each playing a distinct role. Specifically, the abundance of *Ruminococcaceae* correlated with the production of longer-chain acids like butyrate, pentanoate, and hexanoate (ρ: 0.33–0.50). *Marinomonas* and *Psychromonas* were associated with hydrolysis (ρ: 0.33–0.55), while the *Clostridiaceae* population was positively correlated with major acids (e.g., acetic, butyrate acid) (ρ: 0.36–0.52) (Figure 4b; Figure S6).

**Figure 4.**
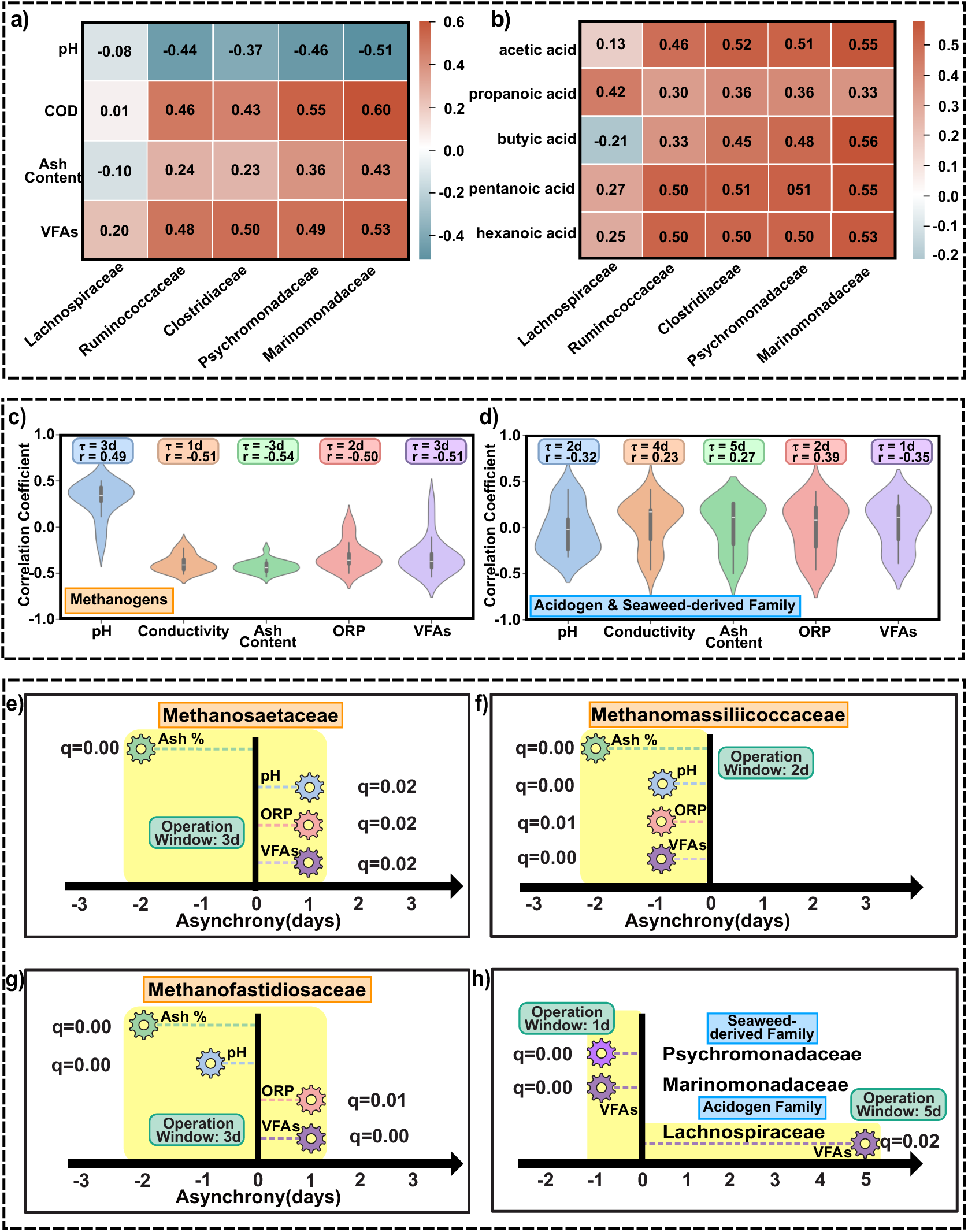
Temporal synchrony analysis and asynchrony-resolved analysis of the two-tier interplay framework. a) Pearson correlation matrix heatmap showing temporal synchrony interplay between engineering parameters and microbial community; b) Pearson correlation matrix correlation heatmap showing temporal synchrony interplay between microbial community and individual VFAs (strong positive links to acetic, propanoic, pentanoic and hexanoic acid). c) Violin plot of asynchrony-resolved EACC analysis of Methanogens and engineering parameters (*Note:* Distributions of ρ(τ) over τ ∈ [−10, +10] d (positive τ: This engineering parameter leads microbe) show a negative correlation with conductivity/ash content/ORP/VFAs and a positive correlation with pH, peaking at short positive τ (∼0–3 d)).; (d) Violin plot of asynchrony-resolved EACC analysis of Acidogens and engineering parameters (*Note:* Distributions of ρ(τ) over τ ∈ [−10, +10] d show a positive correlation with VFAs and EC and a negative correlation with pH, peaking at longer τ (∼1–5 d)); Asynchrony-resolved VAR–Granger results for e) *Methanosaetaceae;* f*) Methanomassiliicoccaceae;* g) *Methanofastidiosaceae*; and h) Seaweed-derived Family (*Psychromonadaceae* and *Marinomonadaceae*) and Acidogen Family (*Lachnospiraceae*) versus engineering parameters (pH, ash content, ORP, VFAs). (*Note:* Colored gears represent control strategy execution; q denotes FDR-adjusted significance; yellow shade indicates the operation window).

Although the temporal synchrony analysis confirmed correlation between two tiers by showing that Tier-1 identifies the dominant acid pools under given redox and salinity states (Figure 4a) and tier-2 pinpoints the specific taxa driving the VFAs production (Figure 4b), it was incapable of resolving the asynchronous inherent to the SW-AAD system characterized by two distinct timescales: rapid changes in engineering parameters (e.g., pH, salinity) and the slow restructuring of the microbial community after shocks (seaweed addition in this study) ^94^.

### A Two-Tier, Asynchrony-Resolved Analysis for Quantifying Two-tier Interplay in SW-AAD

To address the limitations of temporal synchrony analysis and quantify discrepancies between engineering parameters’ and microbial communities’ responses to brown seaweed additions, we conducted an asynchrony-resolved analysis using a two-step approach: event-aligned cross-correlation (EACC) and VAR-Granger tests. (Figure 1d-e, Figure S3). Specifically, the EACC analysis revealed a distinct asynchronous response (Figue 4c–d). The violin plot showed that the decline of methanogen community appeared about 0-3 days after engineering parameters (e.g., pH, ORP, VFA and salinity) changed, at which point their EACC’s correlation coefficient(r) was negatively correlated with conductivity, ash, and VFAs (r: −0.49 to −0.54) and positively correlated with pH (r: ≈0.49) (Figure 4d). This trend aligned with the timescale (2-3 days) in a previous study that the methanogen community composition drastically altered when subjected to the stresses of acidity and high salinity^95^. The analysis also captured the delayed enrichment of acidogen that peaked approximately 1–5 days after engineering parameter change, and the abundance of these acidogen was positively correlated with the VFA and conductivity (r: ≈ 0.23–0.39) (Figure 4d). The EACC analysis explored a two-stage process: a rapid (0–3 day) decline in the methanogen population following the increased acidity and salinity, succeeded by a delayed (1–5 day) acidogen enrichment as the community restructured. This temporal delay was consistent with a previous study showing that the changes in engineering parameters typically preceded microbial community responses by approximately 2–4 days, a delay governed by the fundamental constraints of microbial growth kinetics^96^. The weak statistical correlation (r:-0.35 – 0.39) between the overall acidogen community and engineering parameters reflected the biochemical complexity of acidogenesis^97^, since the production of VFAs is a multi-step process performed by a diverse group of bacteria with varied metabolic pathways and kinetics^98,99^. Therefore, we further applied the stricter method (VAR-Granger test) between two tiers to confirm the asynchrony.

VAR-Granger tests served as confirmation of the directional relationships between the two tiers (Figure 4f-g; Figure S3). For methanogens, changes in pH, ORP, ash content, and VFAs could predict declines of *Methanosaetaceae*, *Methanomassiliicoccaceae*, and *Methanofastidiosaceae* within a 1–3-day (Figure 4e-g). For example, a change in pH preceded a decline in *Methanosaetaceae* by 1 day (False Discovery Rate(q): 0.02) (Figure 4e); a change in ash preceded *Methanomassiliicoccaceae* by 1 day (q: 0.03) (Figure 4f). In the reverse direction, microbial community shifts predicted chemical changes with a much longer time (e.g., *Methanosaetaceae* preceding VFAs by 5 days, q: 0.04) (Figure 4e), which was consistent with the time required for the reactor’s chemistry to adjust after the microbial population had already changed (Figure 4e-g). For acidogen, while the rise in VFAs preceded some acidogens like *Lachnospiraceae* by ∼5 days (q: 0.00) (Figure 4h), the seaweed-derived families *Marinomonadaceae*and *Psychromonadaceae* responded more quickly (within 2-4 days, q: 0.04 – 0.05). They also led to the next increase of VFAs by ∼1 day (q < 0.01) (Figure 4h), acting as the sign for VFAs accumulation. The VAR– Granger results were consistent with the EACC result, indicating the operation window. Collectively, our asynchrony-resolved analysis identifies two patterns in operation window: (1) engineering parameters forecasted microbial shifts, providing a 1–2-day advance of methanogen decline and a 2–5-day forecast of acidogen enrichment (Figure 4e-g), and (2) seaweed-derived microbial communities provided a 1-day forecast of VFA increases (Figure 4h). Quantifying the operational window is critical because it allows the application of control strategies when the system is most favorable for VFAs accumulation, thereby maximizing efficiency.

### Application of Operation Window and Feedback Control Strategy for SW-AAD Systems

Leveraging the asynchronous relationships established between the two tiers (engineering diagnostics and microbial surveillance), feedback control strategies were developed to operate within the operational windows and bolster VFA production in SW-AAD systems. The control strategies simulated in Python consisted of four practical executions for SW-AADs (Table S7, Figure S8): 1) Prolonging HRT. When engineering parameters trended toward threshold (e.g., conductivity ≥ 45 mS cm⁻¹, ash content > 3%, or pH ≤ 5.9 (Figure 2a-c)), the SW-AAD would be operated at an extended HRT in the subsequent three days to prevent the introduction of new substrate and/or alkalinity that could facilitate methanogen recovery. 2) Applying internal circulation. When a high VFA production rate (≥ 350 mg COD L⁻¹ d⁻¹) was observed, or when an increase in the relative abundance of seaweed-derived microbial communities (i.e., *Marinomonadaceae* and *Psychromonadaceae*) was detected, internal circulation would be boosted for a 24-hour period to enhance mixing and improve mass transfer and sustaining robust acidogen activity. 3) Maintaining optimal pH range. The pH in the SW-AAD system would be consistently maintained at a mildly acidic value of 5.6–5.9 (Figure 2a), which would create a constant selective environment favorable for acidogens while suppressing methanogens. 4) Periodic withdrawal of VFAs. If the SW-AAD system exhibits over-acidification (pH < 5.5) or a flattened VFA production rate, ∼10% of the VFA-rich liquor would be withdrew to lower the VFAs content in the system and relieve product inhibition, while preserving the hydraulic and microbial stability, which follows the *in-situ* product removal (ISPR) for acid/product inhibition^100^. Each of these strategies would be executed within the operational window determined from the asynchrony-resolved analysis (Figure 4e-h).

To examine the effectiveness of each feedback control strategy, a simulation was performed using the same input conditions that we used before (Figure 5a, Table S9). The results demonstrated that the feedback control substantially enhanced the VFA production, increasing the cumulative VFA (cVFA) yield by 41.8% over 50 days, from 11.38 to 16.13 g COD L⁻¹ (a net increasing of +4.75 g COD L⁻¹) (Figure 5a). A detailed analysis revealed that most of the improvement (+4.20 g COD L⁻¹) was attributable to Strategy 4 (VFA withdrawal), with smaller contributions from Strategy 1 (Prolonged HRT) (+0.45 g COD L⁻¹) and Strategy 2 (internal circulation) (+0.10 g COD L⁻¹) (Figure 5b). Throughout this AAD phase (after Day 21), The Strategy 3 (pH maintaining acidic) didn’t contribute to cVFA improvement (+0 g COD L⁻¹), however pH was continuously monitored to ensure it remained within the target range of 5.6–5.9. (Figure 5b) The improvement of cVFA indicated that the VFA product inhibition was the primary bottleneck limiting the AAD performance in our benchmark and was consistent with previous studies that *in-situ* product removal boosted VFA productivity ^101^. However, this control model simulation did not resolve complex hydrodynamics or buffer equilibria, since the microbial data was based on relative abundances from a single AAD reactor (Figure 3b-3d), while the operation windows were established from statistical analysis (Figure 4c-4h).

**Figure 5.**
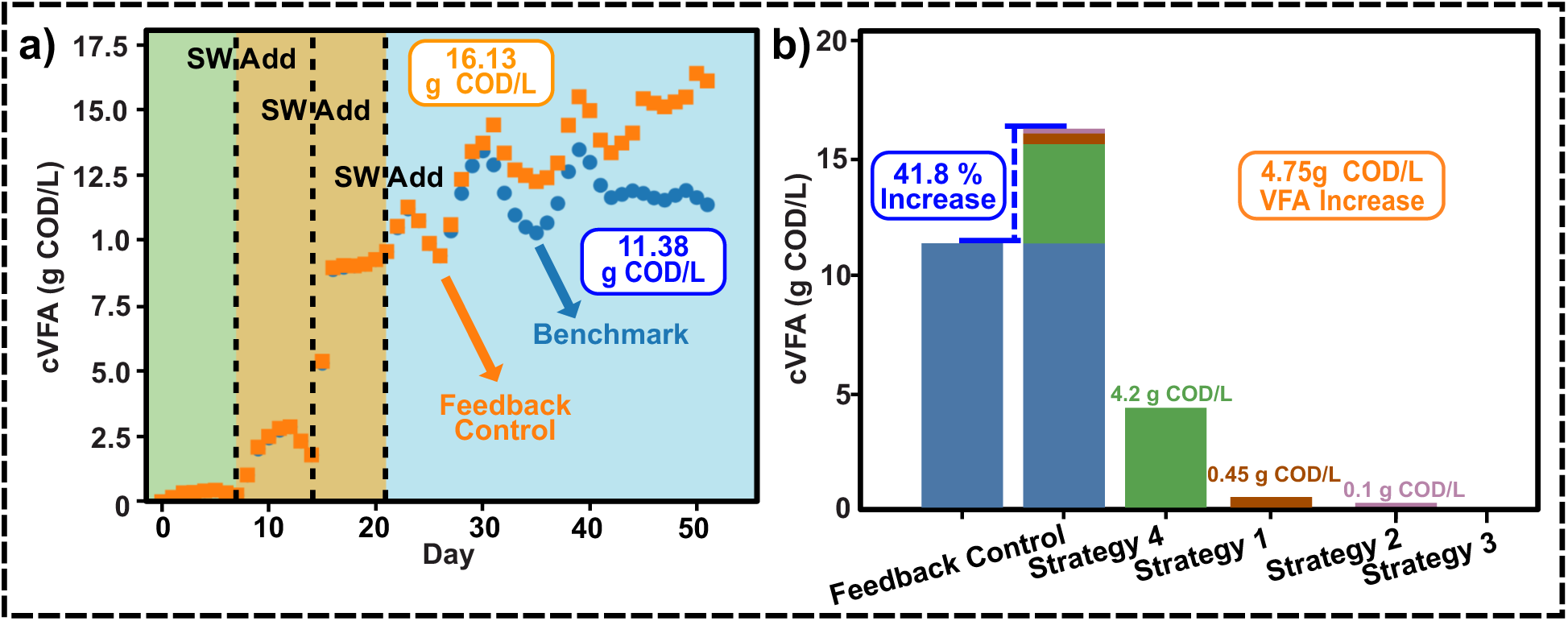
Feedback control model simulation for boosting VFA production in a SW-AAD system. (a) Simulated Cumulative VFAs (cVFA) under the benchmark (without the feedback control) and with feedback control. Daily markers over Days 0–50 with seaweed-addition times (dashed lines). Feedback control leads to higher conversion efficiency, accelerating cVFA from 11.38 to 16.13 g COD·L⁻¹. (b) A detailed analysis of cVFA yield. Feedback total (16.13 g COD·L⁻¹) equals the sum of the benchmark yield (11.38 g COD·L⁻¹), strategy 4 (VFAs withdrawal, 4.20 g COD·L⁻¹), strategy 1 (Prolonged HRT, 0.45 g COD·L⁻¹), strategy 2 (Inner circulation, 0.10 g COD·L⁻¹) and strategy 3(pH maintaining acidic: 0 g COD·L⁻¹). Compared with the benchmark, the feedback control combined with these strategies boosted the VFA yield by 41.8%.

### Comparative Carbon Conversion and Techno-Economic Analysis of Conventional AD versus SW-AAD System

The efficiency of the SW-AAD system was reflected by its carbon conversion efficiency (CCE) (Figure.6a; Table S10), which peaked at 52% from 10% (Figure.6a; Table S10), capturing over half the influent carbon as VFAs. This peak performance represented a 5- to 10-fold greater carbon capture than conventional AD (CCE ≈ 5–10%), establishing the high potential of the SW-AAD platform for valorizing organic waste streams and motivating us to perform the TEA^102^, which compared a SW-AAD system with/without control strategy against a conventional AD with (Table S12) at the same scale (5,600 m^3^, the one that we took digestate samples in GLSD). The SW-AAD system exhibited a substantial capital expenditure (CAPEX) advantage by replacing the AD’s combined heat-and-power (CHP) units with membrane VFAs recovery train. The total CAPEX was reduced by 58% (from $17.9 M to $7.7 M in total) (Figure 6b). However, operational expenditures (OPEX) were estimated as $4.36 million y⁻¹, driven by feedstock and utility costs, compared to $0.54 million y⁻¹ for the conventional AD system that benefited from free municipal sludge and recycled heat. The conventional AD generated $5.06 M in annual revenue ($4.49 M from tipping fees and $0.57 M from electricity) at a cost of $2.65 M, yielding a yearly profit of $2.41 M)(Figure 6b). In contrast, the SW-AAD system with feedback control strategies had an annual yield of 320 tonnes VFAs, valued at $0.69 M. When combined with income from carbon credits ($0.06 M) and tipping (gate) fees ($3.59 M), the gross revenue was $4.34 M (Figure 6b). Factoring in annual expenses of $5.26 M, the AAD system with feedback control strategies would be operated at a $0.92 M deficit, while the AAD system without control yielded a 230 tonnes of VFAs ($0.48 M), resulting in a lower revenue ($4.13 M) and a larger annual deficit of $1.13 M (Table S12).

**Figure 6.**
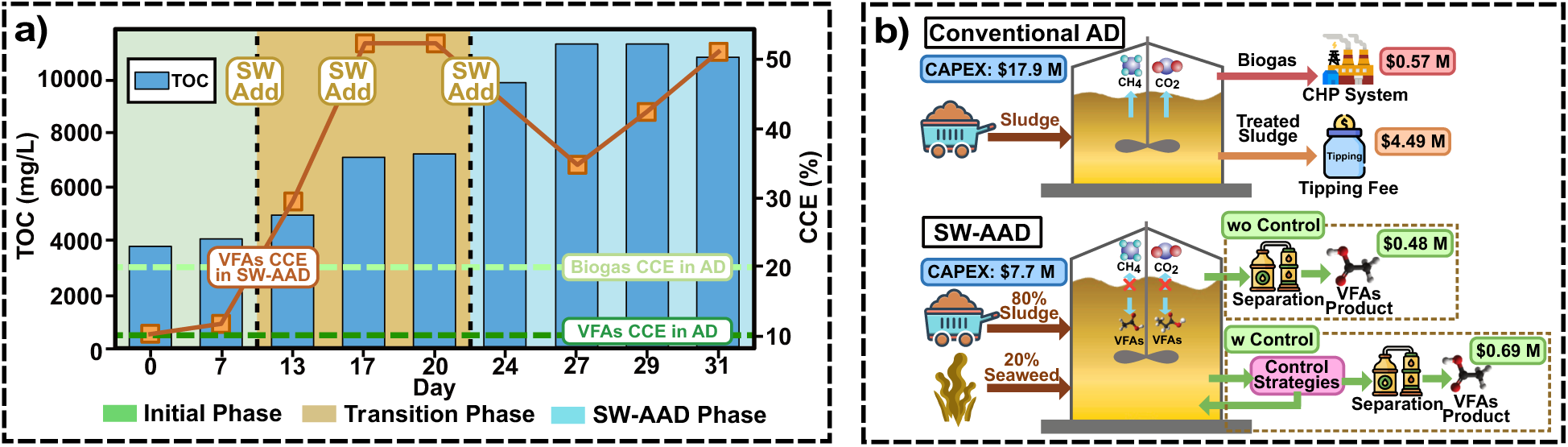
System-level performance of SW-AAD versus conventional AD. (a) Trajectory of total organic carbon (TOC, bars) and carbon-conversion efficiency (CCE, lines). The plot distinguishes three operational windows including start-up (green), transition (amber) and fully implemented SW-AAD (blue), highlighting the step-change in VFA production and CCE once seaweed dosing and electro-saline control are applied; (b) Schematic showing comparison of conventional AD and SW-AAD with/without feedback control.

It should be noted that although the conventional AD system was profitable, its economic model was vulnerable, relying heavily on policy-sensitive tipping fees which constituted over 80% of the total revenue^103^. By contrast, the SW-AAD offers a more resilient alternative through producing high-valued VFAs. Despite this revenue stream, the SW-AAD system’s profitability is highly sensitive to the cost of seaweed feedstock^104^ (Figure S9). We envision that SW-AAD could become economically superior as the value of VFAs augments in matured carbon markets and the regulations become tightened for waste disposal^105^. One feasible pathway leading to the economic viability of SW-AAD is to use negative-cost feedstocks, such as Sargassum^106^, where receiving a tipping fee to manage the problematic biomass can eliminate the primary operational expense and transform SW-AAD into a highly competitive waste-to-value technology.

### Significance of a Two-Tier SW-AAD Framework in Redirecting Organic-Waste Carbon from Biogas to High-Value VFAs

The two-tier SW-AAD platform developed in this study coupled engineering diagnostics (Tier-1, e.g., pH, salinity, VFAs) with microbial surveillance (Tier-2) to resolve the system’s temporal asynchrony. The 70-day experimental results showed that engineering parameters altered before a collapse in the methanogen population by approximately 0–3 days, while acidogenic families rose about 1–5 days later. Two specific seaweed-derived lineages, *Psychromonadaceae* and *Marinomonadaceae*, served as early signals, foreshadowing a next-day VFAs increase. Based on the operation windows obtained from asynchrony-resolved analysis, four feedback control strategies were executed within the operational windows to transform a one-way “inhibited digester” into a time-responsive, controllable VFAs refinery, which could maintain a methane-arrested state and stabilize as a butyrate/hexanoate/acetate pool. Conceptually, this two-tier platform recasts SW-AAD from a static/passive feedstock receiver into an operable biorefinery system where the chemistry sets the state, the operation window signifies execution time, and the control strategies enhance VFA yields.

However, several limitations in this study should be solved in future works. For example, the control thresholds were determined from a lab-scale SW-AAD dataset using interpolated chemistry and weekly generic sequencing data, and the reliance on relative abundances might mask the absolute biomass changes. Furthermore, the VAR-Granger models established asynchrony interplay rather than mechanistic proof. The feedback control strategies were simulated, and the AAD system did not resolve first-principles process dynamics (e.g., ADM1) and was not yet tuned for scale-up hydrodynamics or seasonal feedstock variability. Despite these limitations, this study provides a practical, deployable blueprint for repurposing legacy ADs to achieve higher and more reliable VFA yields using natural biomass. Moreover, the two-tier framework is broadly applicable beyond seaweed to other AAD strategies, enabling a paradigm shift from passive observation to a proactive strategy to “predict, intervene, and monetize” the carbon stream from organic wastes.

## Acknowledge

Microbial Analysis, Resources, and Services, Center for Open Research Resources and Equipment, University of Connecticut.

