## Supporting Information for "Enhancing Volatile Fatty Acid Accumulation in Seaweed-Arrested Anaerobic Digestion via a Two-Tier Framework of Engineering Diagnostics and Microbial Surveillance"

**Test S1 Gravimetric Determination of Salinity via Dry Ashing Method**

Salinity was measured on a wet weight basis using a dry ashing method. Digestate and seaweed samples were analyzed in duplicate. Samples were weighed into pre-weighed ceramic crucibles and placed in a muffle furnace at 80 °C for 2 hours to remove moisture. The temperature was then increased to 600 °C and maintained for 3.5 hours to achieve complete ashing. Crucibles remained covered throughout the process to avoid contamination and sample loss. After cooling to room temperature, crucibles were reweighed. Salinity was calculated as the percentage of ash relative to the initial wet weight of the sample. This procedure was adapted from the protocol established by Zhang et al. (2025), which was originally developed based on AOAC Official Methods 942.05 for the determination of ash content[1]. The mass of the empty crucible was recorded as $W_0$. After loading the wet sample, the combined mass was recorded as $W_1$. After ashing at 600 °C, the final mass corresponding to the crucible and remaining inorganic residue was recorded as $W_2$. The ash content was calculated as follows:

$$\text{Ash content} = \frac{w_2 - w_0}{w_1 - w_0}$$

**Test S2 Gas Chromatography–Mass Spectrometry (GC-MS) Profiling of VFAs in SW-AAD System**

VFAs were first extracted using a liquid–liquid extraction (LLE) method (Test S3). The organic phase was then analyzed by gas chromatography–mass spectrometry (GC–MS) using an Agilent 7820A GC coupled with a 5975 MSD (Agilent Technologies, USA).Separation was carried out on a DB-FFAP capillary column (25 m × 0.32 mm × 0.5 µm; Agilent Technologies). Helium was employed as the carrier gas at a constant flow rate of 1.0 mL/min. The GC oven was initially held at 80 °C for 2 minutes, then ramped at 10 °C/min to a final temperature of 230 °C, which was maintained for an additional 2 minutes. The total run time was 21 minutes[2].Samples (1 μL) were injected in split mode (50:1) with the injector temperature set at 250 °C. The ion source and interface temperatures were maintained at 230 °C and 220 °C, respectively. A solvent cut time of 2 minutes was applied to eliminate solvent interference during detection. The mass spectrometer was operated in full scan acquisition mode. Sample introduction was performed using an automated liquid sampler to ensure injection consistency. Calibration curves were constructed using a stepwise concentration gradient, as detailed in Text S4.

**Test S3 Separation of VFAs from Aqueous Phase via Liquid–Liquid Extraction (LLE)**

The sample was initially diluted by transferring 1 mL of the original liquid matrix into a 15 mL centrifuge tube, followed by the addition of 9 mL deionized water to achieve a 10-fold dilution, ensuring that analyte concentrations fell within the optimal detection range. The diluted samples were then centrifuged at 2,600 rpm for 10 minutes to remove particulate matter. The resulting supernatant was carefully filtered using a 10 μm syringe filter into clean vials; syringe filters were replaced after every 2–3 uses to prevent clogging and cross-contamination[3].

The filtered solution was acidified by adjusting the pH to between 2 and 3 through the dropwise addition of 90% nitric acid, facilitating the protonation of VFAs and enhancing their extractability. For the LLE procedure, 5 mL of the acidified sample was transferred into a clean separatory funnel, and an equal volume of methyl tert-butyl ether (MTBE) (HPLC grade, Sigma) was added. The mixture was then shaken vigorously for 5 minutes to promote efficient partitioning of VFAs into the organic phase. After phase separation, the bottom aqueous layer was discarded, and the upper organic layer containing the extracted VFAs was collected for subsequent GC-MS analysis[4].

**Test S4 VFAS Standard Calibration Curve by GC-MS**

Ten individual VFAs standards, including formic acid, acetic acid, propanoic acid, isobutyric acid, butanoic acid, isovaleric acid, pentanoic acid, isocrotonic acid, hexanoic acid, and heptanoic acid, were each dissolved in methyl tert-butyl ether (MTBE) to prepare stock solutions at a concentration of 50 mM. A series of two-fold serial dilutions was subsequently performed to generate a calibration range extending down to 0.1953 mM. External calibration curves constructed from these standards exhibited excellent linearity, with all correlation coefficients ($R^2$) greater than 0.98. No internal standard was used in the analysis. Compound identification was based on retention time and diagnostic mass-to-charge (m/z) ratios under full scan mode. The retention times and corresponding m/z values were as follows: formic acid (retention time = 6.822 min, m/z 29, 45), acetic acid (retention time = 6.107 min, m/z 43, 60), propanoic acid (retention time = 7.182 min, m/z 74, 45, 28), isobutyric acid (retention time = 7.510 min, m/z 73), butanoic acid (retention time = 8.262 min, m/z 60, 73), isovaleric acid (retention time = 8.732 min, m/z 60, 87), pentanoic acid (retention time = 9.539 min, m/z 60, 73), isocrotonic acid (retention time = 10.248 min, m/z 57, 73), hexanoic acid (retention time = 10.717 min, m/z 60, 73), and heptanoic acid (retention time = 11.852 min, m/z 60, 73, 87).

**Test S5 Data processing by Shape-preserving monotone cubic interpolation (PCHIP)**

To facilitate a direct comparison between the engineering parameters and the microbial surveillance data, which were collected at different frequencies, it was necessary to harmonize all time-series to a uniform daily grid (Day 0–70). This was achieved using the PCHIP method. Unlike standard spline interpolation, which can introduce artificial oscillations or "overshoots" between measured data points, PCHIP is specifically designed to be shape-preserving. This means that the interpolated values will always respect the monotonicity of the original data; if the measured data points are increasing, the interpolated curve between them will also be increasing. This property is critical for preserving the integrity of the underlying physical and biological trends in the reactor, ensuring that the subsequent correlation and asynchrony analyses were performed on data that accurately reflected the system's dynamics.

### Text S6 Interplay Analysis via Pearson cross correlations matrix

To establish a baseline understanding of the simultaneous relationships between the reactor's physical-chemical state and its microbial community, we performed a **Pearson cross correlations matrix**. Pearson's correlation coefficient (*r*) measures the strength and direction of a linear relationship between two continuous variables, with values ranging from -1 (perfect negative correlation) to +1 (perfect positive correlation). In this context, "cross-domain" signifies that we computed correlations between variables from the two different tiers of our framework: the engineering diagnostics and the microbial surveillance. This analysis was comprehensive and included the following specific comparisons: **Key Engineering Parameters vs. Microbial Guilds:** The relationships between pH, COD, ash content, and total VFAs were tested against the relative abundances of the methanogen and acidogen guilds. **Biogas Production vs. Methanogens:** The production of $CH_4$ and $CO_2$ was correlated specifically with the abundance of methanogenic families. **VFA Speciation vs. Acidogenic Families:** The concentrations of individual VFA species (acetate, propionate, butyrate, pentanoate, hexanoate) were correlated with the relative abundances of key acidogenic families to link specific products to potential producers. This analysis provided a static "snapshot" of the system's interplay at matched time points, identifying which engineering parameters and microbial taxa were strongly associated.

**Text S7 Quantifying Two-tier Interplay via Asynchrony-resolved Analysis**

Our analytical strategy employs a two-step method that combines Event-Aligned Cross-Correlation (EACC) with Vector Autoregression (VAR) Granger Causality testing. EACC is to first explore and identify significant temporal relationships, then VAR-Granger test formally test them for directional, predictive causality. EACCfocused on data windows centered around each of the three seaweed-addition events. Within these windows, we compared "two-tier pairs"—each consisting of one engineering time-series (e.g., daily pH) and one microbial time-series (e.g., daily relative abundance of *Methanosaetaceae*).The cross-correlation was computed for each pair by systematically shifting one time-series relative to the other across a range of integer time asynchrony ($\tau$) from -10 to +10 days. A positive $\tau$ signifies that the engineering parameter leads the microbial response. From this analysis, we extracted two key metrics for each event: the peak correlation magnitude and its corresponding optimal time $\tau$, which represents the time delay with the strongest statistical relationship. The results of this analysis are summarized in Table S4.

To verify the direction for each two-tier pair, we used bivariate vector autoregression (VAR)–based Granger causality tests. This process involved the following steps: **Data Preprocessing:** Within each event window, time-series data were mean-centered by subtracting the window's mean from each value. This removes level offsets without altering the timing of the signals. **Model Selection:** The VAR model asynchrony order ($p$), ranging from 1 to 5 days, was selected by minimizing the Akaike Information Criterion (AIC). This criterion balances model goodness-of-fit with model complexity to prevent overfitting. **Hypothesis Testing:** We tested for predictive relationships in both directions (from parameter to microbe and from microbe to parameter) using F-tests on the lagged terms of the VAR model. **Statistical Correction:** To account for the large number of comparisons, the familywise error rate was controlled using a False Discovery Rate (FDR) adjustment, with a significance threshold of $q < 0.05$. For all significant predictive links, we report the asynchrony time (in days) and the associated q-value. This converts statistical timing patterns into the directional links used for feedback control. The full per-pair outputs appear in Tables S5 and S6.

#### Text S8: Derivation of the Feedback Control Protocol

The feedback control strategy was designed to convert the statistical findings from the asynchrony analysis into a set of actionable, data-driven rules. The specific trigger thresholds were derived from the empirical distributions observed during the 70-day experiment to ensure they were representative of the system's dynamics. The detailed rules are presented in Table S7, and the methodology for their derivation is as follows:

- **Trigger Thresholds:** Thresholds were set using statistical percentiles of the relevant time-series data to remain data-driven yet generalizable.
    - The trigger for a "rising" VFAs accumulation rate was set at the 60th percentile of positive daily VFA changes.
    - The "flattening peak" condition for VFA draw-off was defined using the 25th percentile of the absolute rate of change on high-VFA days, with the VFA concentration level set at the 75th percentile.
    - Salinity and redox shock indicators were based on the 75th percentile of conductivity and ORP values observed immediately after seaweed dosing.
    - The operational pH band was defined by the 25th and 75th percentiles of pH values observed during periods of high VFA concentration.

- Microbial triggers for seaweed-derived taxa (*Marinomonadaceae*, *Psychromonadaceae*) were set at the 75th percentile of positive day-over-day changes.

- **Controller Stability:** To prevent rapid, unstable switching of control actions (i.e., "controller chattering"), we implemented two stability measures: **Minimum Dwell Times:** Once an action was triggered, it was held for a minimum duration (e.g., 48–72 hours for an HRT hold, 12–18 hours for a circulation pulse). **Hysteresis:** A ±5% hysteresis band was applied to all numerical comparators, meaning the value must fall significantly below a threshold before the control action is turned off.

### Text S9: TEA Method

To ensure a fair comparison, the annualized cost of each system is calculated using equations 1-2.

$$\text{Annualized cost } = CRF \times CAPEX + Opex \quad (1)$$

$$\text{CRF} = \frac{\mathrm{i}(1+\mathrm{i})^{\mathrm{n}}}{(1+i)^{n}-1} \quad (2)$$

The capital recovery factor (CRF) is used to convert the capital expenditure (CAPEX) into equal annual payments over the project lifetime and operational expenditure (OPEX) is the annual operation cost. For the capital expenditure assessment of the anaerobic digestion biogas facility, we utilized reported numbers from published studies. Studies indicates that the establishment of a standard anaerobic digestion (AD) biogas facility costs expenditures of roughly $200 per ton of feedstock, with the digester comprising merely 20–30% of the overall capital expenditure (CAPEX), while the combined heat and power (CHP) unit constitutes the predominant portion of the investment (50–60%)[5]. To estimate the CAPEX for the VFA manufacturing units, we used the conventional scaling factor method to calculate the cost of our system based on the reported values from Equation 4 shows how we calculate the CPAEX cost[6]. Based on the literature[7], the VFA recovery and separation units usually make up about 40% of the overall CAPEX, while the AD system is responsible for the other 60% of the cost when it comes to producing VFA. The approach for estimating CAPEX is further detailed in the Supplementary Material. The OPEX for each system was determined based on its corresponding CAPEX. For the SW-AAD system, we incorporated an extra expense for seaweed feedstock, estimated at $200 per ton. Finally, we computed the revenue and profit produced by each system using Equation 5.

$$\text{Annual Profit} = \text{Revenue} - \text{Annualized cost} \tag{5}$$

Revenue sources are the selling of power for the traditional AD system and volatile fatty acids for the SW-AAD system. Moreover, various studies underscore the significance of tipping fees (or gate fees) imposed by waste management facilities for the acceptance of waste. These fees constitute an essential revenue stream, mitigating significant operational costs including personnel, maintenance, and regulatory compliance. In several U.S. facilities, tipping costs may surpass $100 per ton, facilitating everyday operations and financing recycling initiatives and infrastructure enhancements[8]. A tipping cost of $50 per ton is used for our analysis.

**Table S1.** Comprehensive Engineering Parameter Monitoring

| Date | Day | Sample Component | NH4+,mg/L | pH | Conductivity, ms/cm | COD, mg/L | ORP, mV |
|---|---|---|---|---|---|---|---|
| 9/11/2024 | 1 | GLSD Digestate | 1540 | 7.83 | 11.65 | 19000 | -239 |
| 9/13/2024 | 3 | | 1580 | 7.84 | 10.34 | 15200 | -245 |
| 9/16/2024 | 6 | | 1600 | 7.86 | 11.04 | 16400 | -269 |
| 9/18/2024 | 8 | | 1670 | 7.78 | 10.48 | 15400 | -233 |
| 9/20/2024 | 10 | 50% Digestate + 50% Seaweed | 895 | 7.43 | 17.8 | 15600 | -221 |
| 9/23/2024 | 13 | | 1040 | 7.45 | 24.6 | 24200 | -251 |
| 9/25/2024 | 15 | | 1150 | 7.5 | 27.6 | 21800 | -227 |
| 9/27/2024 | 17 | 25% Digestate + 75% Seaweed | 734 | 6.57 | 35.6 | 29300 | -182 |
| 9/30/2024 | 20 | | 629 | 6.39 | 38.6 | 32600 | -210 |
| 10/2/2024 | 22 | | 613 | 6.26 | 39.7 | 33100 | -189 |
| 10/4/2024 | 24 | 12.5% Digestate + 87.5% Seaweed | 451 | 5.98 | 40.2 | 33820 | -132 |
| 10/7/2024 | 27 | | 360 | 5.68 | 41.2 | 35400 | -122 |
| 10/9/2024 | 29 | | 374 | 5.64 | 42.6 | 41000 | -123 |
| 10/11/2024 | 31 | | 384 | 5.67 | 42.8 | 42400 | -143 |
| 10/14/2024 | 34 | | 298 | 5.65 | 43.5 | 47600 | -112 |
| 10/16/2024 | 36 | | 353 | 5.74 | 43,2 | 43200 | -153 |
| 10/18/2024 | 38 | | 365 | 5.72 | 44.9 | 44400 | -133 |
| 10/21/2024 | 40 | | 414 | 5.93 | 44.6 | 41200 | -140 |
| 10/23/2024 | 43 | | 614 | 5.7 | 44.7 | 46000 | -121 |
| 10/25/2024 | 45 | | 488 | 5.74 | 44.7 | 43300 | -123 |
| 10/28/2024 | 48 | | 532 | 5.83 | 47.4 | 39920 | -108 |
| 10/30/2024 | 50 | | 459 | 6.02 | 47.5 | 42900 | -106 |
| 11/1/2024 | 52 | | 519 | 5.84 | 47.6 | 44800 | -188 |
| 11/4/2024 | 55 | | 380 | 5.9 | 43.8 | 42100 | -139 |
| 11/6/2024 | 57 | | 580 | 5.87 | 47.4 | 45600 | -133 |
| 11/8/2024 | 59 | | 580 | 6.03 | 43.4 | 41200 | -146 |
| 11/11/2024 | 62 | | 391 | 5.94 | 48.8 | 39800 | -179 |
| 11/13/2024 | 64 | | 516 | 6.31 | 42.1 | 37300 | -150 |
| 11/15/2024 | 66 | | 598 | 5.9 | 44.4 | 31700 | -142 |
| 11/18/2024 | 68 | | 462 | 6.09 | 45.6 | 35300 | -132 |
| 11/20/2024 | 71 | | 492 | 5.87 | 47.2 | 34200 | -115 |

**Table S2** Comparison of VFA yield and key microbial change with other AAD research

| Inhibition method | Target / pathway | VFA yield / profile | Key microbial changes |
|---|---|---|---|
| 2-Bromoethanesulfonate (BES)[9] | Coenzyme-M reductase (methanogens) | 49.3 g $L^{-1}$ (Ac/Pr/Bu) | Methanogens drop; Clostridia rise |
| Hydrogen peroxide ($H_2O_2$)[10] | Oxidative damage to methanogens | 1.23 g $L^{-1}$ (Ac & Pr) | Archaea inactivated (not specified) |
| High pH (~9)[11] | Alkalinity stress on all methanogens | 80.6 g COD $L^{-1}$ (≈50 g VFA $L^{-1}$) | Alkaliphilic fermenters dominate (e.g. Anaerosalibacter) |
| Low pH + heat pretreatment[12] | Acid shock + thermal kill of methanogens | 78 g $L^{-1}$ total VFA | Spore-forming Clostridia enriched |
| Micro-aeration (initial $O_2$)[13] | $O_2$ toxicity to strict anaerobes | 0.80 g VFA $g^{-1}$ VS | Methanogens suppressed; facultative fermenters rise |
| Thermal shock (70–100 °C)[14] | Heat kill of non-spore methanogens | 37 g $L^{-1}$ (Ac & Bu) | Spore-forming fermenters dominate |
| Bioaugmentation + BES[15] | Block methanogens & enrich homoacetogens | 49.3 g $L^{-1}$ acetate (2.19 g $g^{-1}$ VS) | Acetobacterium & acetogens dominate; methanogens drop |
| Alkaline + thermophilic fermentation (pH 10, 55 °C)[16] | Thermal and pH stress on methanogens | 4,748 mg COD/L | Methanogens suppressed; acidogenesis enhanced |
| Ultrasonic + alkaline (pH 10)[17] | Cell lysis + methanogen inhibition via pH | Enhanced (not specified) | Enhanced hydrolysis; increased soluble COD |
| Alkaline fermentation (pH 10)[18] | Methanogen inhibition via alkaline stress | 1,721 mg COD/L | Clostridia rise; Methanogenic archaea drop |
| 50 mM 2-BES[19] | Methanogenesis enzyme inhibition | 7,266 mg COD/L | Methanogens drop; fermentative DOM rise; denitrification improved |

**Table S3** Comparison between EACC and VAR-Granger Test

| | Event-Aligned Cross-Correlation | VAR-Granger Test |
|---|---|---|
| **Goal** | Respond Pattern | Predictive Direction |
| **Relationship Type** | Correlation & Time Lag | Predictive Causality / Information Flow |
| **Data Requirement** | Requires specific, repeating "events" to align the data. | Dstable statistical properties over time |
| **Method** | Signal processing & Averaging | Regression model comparison & Hypothesis testing |
| **Advantage** | Noise Reduction; Good for Event-Driven Systems | Multivariate Analysis， Statistically Rigorous |
| **Disadvantage** | Shows Correlation, Not Causality; Relatively Simplistic | Strict Data Requirements； Sensitive to Model Parameters |

**Table S4** PCHIP-type interpulated daily parameter and microbial data

| Day | pH | Conductivity, ms/cm | Ash Content, % | ORP, mV | COD/NH4 | VFAs, mg COD/L | Psychromonadaceae, | Marinomonadaceae |
|---|---|---|---|---|---|---|---|---|
| 0 | 7.83 | 11.65 | 0.40 | -239.0 | 12.34 | 1037 | 0.00 | 0.00 |
| 1 | 7.83 | 10.74 | 0.40 | -241.2 | 10.48 | 1201 | 0.72 | 0.30 |
| 2 | 7.84 | 10.34 | 0.40 | -245.0 | 9.62 | 1317 | 1.07 | 0.45 |
| 3 | 7.85 | 10.52 | 0.40 | -253.1 | 9.78 | 1392 | 1.07 | 0.45 |
| 4 | 7.86 | 10.86 | 0.41 | -263.7 | 10.09 | 1450 | 1.07 | 0.45 |
| 5 | 7.86 | 11.04 | 0.41 | -269.0 | 10.25 | 1473 | 1.07 | 0.45 |
| 6 | 7.84 | 10.76 | 0.41 | -253.3 | 9.74 | 1378 | 0.54 | 0.22 |
| 7 | 7.78 | 10.48 | 0.41 | -233.0 | 9.22 | 1282 | 0.00 | 0.00 |
| 8 | 7.59 | 13.43 | 0.61 | -224.8 | 12.65 | 2063 | 0.53 | 0.23 |
| 9 | 7.43 | 17.80 | 0.94 | -221.0 | 17.43 | 3064 | 1.07 | 0.45 |
| 10 | 7.43 | 20.43 | 1.25 | -228.8 | 20.15 | 3478 | 1.07 | 0.45 |
| 11 | 7.44 | 22.68 | 1.56 | -243.2 | 22.36 | 3783 | 1.07 | 0.45 |
| 12 | 7.45 | 24.60 | 1.74 | -251.0 | 23.27 | 3903 | 1.07 | 0.45 |
| 13 | 7.48 | 26.00 | 1.80 | -242.9 | 21.12 | 3366 | 1.47 | 0.37 |
| 14 | 7.50 | 27.60 | 1.84 | -227.0 | 18.96 | 2828 | 1.87 | 0.28 |
| 15 | 7.06 | 31.73 | 1.89 | -200.6 | 27.96 | 6349 | 1.47 | 0.37 |
| 16 | 6.57 | 35.60 | 1.95 | -182.0 | 39.92 | 9939 | 1.07 | 0.45 |
| 17 | 6.49 | 36.96 | 2.00 | -189.3 | 45.28 | 10027 | 1.07 | 0.45 |
| 18 | 6.44 | 37.88 | 2.06 | -202.7 | 49.33 | 10067 | 1.07 | 0.45 |
| 19 | 6.39 | 38.60 | 2.11 | -210.0 | 51.83 | 10135 | 1.07 | 0.45 |
| 20 | 6.33 | 39.24 | 2.16 | -203.3 | 52.83 | 10311 | 2.88 | 1.23 |
| 21 | 6.26 | 39.70 | 2.21 | -189.0 | 54.00 | 10621 | 4.69 | 2.00 |
| 22 | 6.13 | 39.97 | 2.30 | -158.2 | 62.73 | 11534 | 2.88 | 1.22 |
| 23 | 5.98 | 40.20 | 2.41 | -132.0 | 74.99 | 12260 | 1.07 | 0.45 |
| 24 | 5.86 | 40.48 | 2.52 | -126.6 | 83.61 | 11793 | 1.07 | 0.45 |
| 25 | 5.75 | 40.80 | 2.62 | -123.2 | 91.41 | 10925 | 1.07 | 0.45 |
| 26 | 5.68 | 41.20 | 2.72 | -122.0 | 98.33 | 10458 | 1.07 | 0.45 |
| 27 | 5.65 | 41.97 | 2.80 | -122.3 | 105.41 | 11410 | 4.56 | 2.67 |
| 28 | 5.64 | 42.60 | 2.87 | -123.0 | 109.63 | 12847 | 8.05 | 4.89 |
| 29 | 5.65 | 42.71 | 2.94 | -133.2 | 110.03 | 13904 | 4.56 | 2.67 |
| 30 | 5.67 | 42.80 | 3.01 | -143.0 | 110.42 | 14477 | 1.07 | 0.45 |
| 31 | 5.66 | 43.04 | 3.10 | -135.0 | 123.53 | 13946 | 1.07 | 0.45 |
| 32 | 5.66 | 43.35 | 3.18 | -120.0 | 147.11 | 12871 | 1.07 | 0.45 |
| 33 | 5.65 | 43.50 | 3.23 | -112.0 | 159.73 | 12026 | 1.07 | 0.45 |
| 34 | 5.70 | 43.35 | 3.24 | -132.5 | 141.24 | 11565 | 1.07 | 0.45 |
| 35 | 5.74 | 43.20 | 3.25 | -153.0 | 122.38 | 11339 | 1.07 | 0.45 |
| 36 | 5.73 | 44.05 | 3.26 | -143.0 | 122.01 | 11723 | 5.42 | 3.95 |
| 37 | 5.72 | 44.90 | 3.26 | -133.0 | 121.64 | 12473 | 9.77 | 7.44 |
| 38 | 5.82 | 44.75 | 3.27 | -136.5 | 112.78 | 13682 | 5.42 | 3.95 |
| 39 | 5.93 | 44.60 | 3.28 | -140.0 | 99.52 | 14526 | 1.07 | 0.45 |
| 40 | 5.87 | 44.63 | 3.29 | -135.1 | 88.91 | 14049 | 1.07 | 0.45 |
| 41 | 5.76 | 44.67 | 3.29 | -125.9 | 79.18 | 13163 | 1.07 | 0.45 |
| 42 | 5.70 | 44.70 | 3.30 | -121.0 | 74.92 | 12686 | 1.07 | 0.45 |
| 43 | 5.71 | 44.70 | 3.31 | -122.0 | 81.83 | 12816 | 3.89 | 2.59 |
| 44 | 5.74 | 44.70 | 3.32 | -123.0 | 88.73 | 12946 | 6.71 | 4.73 |
| 45 | 5.76 | 45.38 | 3.33 | -119.5 | 85.18 | 12854 | 5.25 | 3.62 |
| 46 | 5.79 | 46.66 | 3.33 | -112.6 | 78.59 | 12683 | 2.53 | 1.56 |
| 47 | 5.83 | 47.40 | 3.34 | -108.0 | 75.04 | 12591 | 1.07 | 0.45 |
| 48 | 5.94 | 47.46 | 3.34 | -106.6 | 84.25 | 12779 | 1.07 | 0.45 |
| 49 | 6.02 | 47.50 | 3.35 | -106.0 | 93.46 | 12967 | 1.07 | 0.45 |
| 50 | 5.93 | 47.56 | 3.36 | -147.0 | 89.89 | 12690 | 3.13 | 2.42 |
| 51 | 5.84 | 47.60 | 3.37 | -188.0 | 86.32 | 12413 | 5.18 | 4.39 |
| 52 | 5.86 | 46.61 | 3.37 | -176.4 | 92.66 | 12811 | 4.11 | 3.37 |
| 53 | 5.88 | 44.79 | 3.37 | -153.9 | 104.45 | 13551 | 2.14 | 1.47 |
| 54 | 5.90 | 43.80 | 3.37 | -139.0 | 110.79 | 13949 | 1.07 | 0.45 |
| 55 | 5.89 | 45.60 | 3.37 | -134.8 | 96.24 | 13396 | 1.07 | 0.45 |
| 56 | 5.87 | 47.40 | 3.37 | -133.0 | 78.62 | 12843 | 1.07 | 0.45 |
| 57 | 5.95 | 45.40 | 3.37 | -137.5 | 73.29 | 12926 | 1.29 | 0.94 |
| 58 | 6.03 | 43.40 | 3.37 | -146.0 | 71.03 | 13008 | 1.51 | 1.43 |
| 59 | 6.01 | 44.80 | 3.37 | -158.1 | 79.00 | 12973 | 1.40 | 1.18 |
| 60 | 5.96 | 47.40 | 3.37 | -172.2 | 93.82 | 12875 | 1.18 | 0.70 |
| 61 | 5.94 | 48.80 | 3.37 | -179.0 | 101.79 | 12722 | 1.07 | 0.45 |
| 62 | 6.13 | 45.45 | 3.37 | -166.1 | 89.95 | 11955 | 1.07 | 0.45 |
| 63 | 6.31 | 42.10 | 3.37 | -150.0 | 72.29 | 11276 | 1.07 | 0.45 |
| 64 | 6.11 | 43.05 | 3.37 | -145.5 | 59.74 | 11372 | 1.07 | 0.44 |
| 65 | 5.90 | 44.40 | 3.37 | -142.0 | 53.01 | 11467 | 1.07 | 0.42 |
| 66 | 6.00 | 45.06 | 3.37 | -137.2 | 64.71 | 10922 | 1.07 | 0.43 |
| 67 | 6.09 | 45.60 | 3.37 | -132.0 | 76.41 | 10376 | 1.07 | 0.45 |
| 68 | 6.07 | 46.16 | 3.37 | -126.6 | 76.15 | 10430 | 1.04 | 0.48 |
| 69 | 6.00 | 46.69 | 3.36 | -120.9 | 74.37 | 10683 | 0.95 | 0.53 |
| 70 | 5.87 | 47.20 | 3.35 | -115.0 | 69.51 | 11274 | 0.83 | 0.59 |

**Table S5** Cross-Correlation result between parameters with Methoanogen and Acidogen

| | pH ↔ Methanogen | pH ↔ Acidogen | Conductivity↔ Methanogen | Conductivity ↔ Acidogen | Ash Content ↔ Methanogen | Ash Content ↔ Acidogen | ORP ↔ Methanogen | ORP ↔ Acidogen | VFAs ↔ Methanogen | VFAs ↔ Acidogen |
|---|---|---|---|---|---|---|---|---|---|---|
| **-10.00** | 0.46 | 0.39 | -0.49 | -0.32 | -0.48 | -0.32 | -0.49 | -0.43 | -0.44 | -0.38 |
| **-9.00** | 0.43 | 0.41 | -0.49 | -0.46 | -0.48 | -0.50 | -0.38 | -0.32 | -0.45 | -0.39 |
| **-8.00** | 0.35 | 0.34 | -0.38 | -0.39 | -0.38 | -0.41 | -0.27 | -0.32 | -0.36 | -0.36 |
| **-7.00** | 0.34 | 0.33 | -0.26 | -0.31 | -0.24 | -0.31 | -0.43 | -0.46 | -0.33 | -0.32 |
| **-6.00** | 0.50 | 0.31 | -0.43 | -0.29 | -0.42 | -0.29 | -0.39 | -0.21 | -0.53 | -0.31 |
| **-5.00** | 0.34 | 0.09 | -0.35 | -0.13 | -0.41 | -0.18 | -0.32 | -0.04 | -0.31 | -0.13 |
| **-4.00** | 0.31 | 0.07 | -0.32 | -0.04 | -0.39 | -0.06 | -0.43 | -0.20 | -0.29 | -0.08 |
| **-3.00** | 0.45 | 0.04 | -0.49 | 0.03 | -0.54 | -0.01 | -0.39 | -0.11 | -0.46 | -0.04 |
| **-2.00** | 0.38 | -0.14 | -0.46 | 0.12 | -0.48 | 0.03 | -0.29 | 0.20 | -0.39 | 0.17 |
| **-1.00** | 0.34 | -0.20 | -0.39 | 0.15 | -0.38 | 0.08 | -0.40 | 0.08 | -0.39 | 0.11 |
| **0.00** | 0.48 | -0.24 | -0.51 | 0.19 | -0.50 | 0.11 | -0.36 | 0.12 | -0.51 | 0.21 |
| **1.00** | 0.36 | -0.25 | -0.45 | 0.18 | -0.47 | 0.14 | -0.32 | 0.24 | -0.33 | 0.35 |
| **2.00** | 0.40 | -0.32 | -0.41 | 0.19 | -0.40 | 0.19 | -0.50 | 0.39 | -0.41 | 0.17 |
| **3.00** | 0.49 | -0.32 | -0.47 | 0.19 | -0.46 | 0.21 | -0.45 | 0.19 | -0.54 | 0.23 |
| **4.00** | 0.25 | -0.29 | -0.39 | 0.23 | -0.50 | 0.23 | -0.17 | 0.22 | -0.23 | 0.32 |
| **5.00** | 0.28 | -0.19 | -0.39 | 0.22 | -0.44 | 0.27 | -0.39 | 0.35 | -0.26 | 0.27 |
| **6.00** | 0.30 | -0.27 | -0.42 | 0.19 | -0.45 | 0.27 | -0.30 | -0.02 | -0.37 | 0.11 |
| **7.00** | 0.11 | -0.24 | -0.45 | 0.20 | -0.44 | 0.27 | -0.01 | 0.12 | 0.10 | 0.25 |
| **8.00** | -0.10 | 0.03 | -0.27 | 0.33 | -0.35 | 0.27 | 0.07 | 0.22 | -0.37 | 0.35 |
| **9.00** | 0.02 | 0.03 | -0.23 | 0.17 | -0.37 | 0.26 | -0.06 | 0.27 | -0.11 | 0.15 |
| **10.00** | -0.22 | -0.02 | -0.18 | 0.41 | -0.40 | 0.27 | -0.31 | -0.46 | 0.30 | 0.00 |

**Table S6** Granger Causality p values for parameter and bacteria

| **Methanogen** | | | | | | | | | | |
|---|---|---|---|---|---|---|---|---|---|---|
| **Lag** | **pH→Methanogen** | **Conductivity, ms/cm→Methanogen** | **Ash Content, %→Methanogen** | **ORP, mV→Methanogen** | **VFAs, mg COD/L→Methanogen** | **Methanogen→pH** | **Methanogen→Conductivity, ms/cm** | **Methanogen→Ash Content, %** | **Methanogen→ORP, mV** | **Methanogen→VFAs, mg COD/L** |
| **Lag 1** | 0.12 | 0.43 | 0.61 | 0.69 | 0.03 | 0.02 | 0.01 | 0.01 | 0.01 | 0.01 |
| **Lag 2** | 0.16 | 0.17 | 0.01 | 0.10 | 0.04 | 0.01 | 0.00 | 0.00 | 0.02 | 0.00 |
| **Lag 3** | 0.07 | 0.06 | 0.00 | 0.29 | 0.05 | 0.30 | 0.03 | 0.02 | 0.38 | 0.08 |
| **Lag 4** | 0.08 | 0.42 | 0.00 | 0.31 | 0.13 | 0.80 | 0.61 | 0.00 | 0.38 | 0.72 |
| **Lag 5** | 0.02 | 0.77 | 0.00 | 0.25 | 0.00 | 0.85 | 0.64 | 0.00 | 0.52 | 0.74 |
| **Acidogen** | | | | | | | | | | |
| **Lag** | **pH→Acidogen** | **Conductivity, ms/cm→Acidogen** | **Ash Content, %→Acidogen** | **ORP, mV→Acidogen** | **VFAs, mg COD/L→Acidogen** | **Acidogen→pH** | **Acidogen→Conductivity, ms/cm** | **Acidogen→Ash Content, %** | **Acidogen→ORP, mV** | **Acidogen→VFAs, mg COD/L** |
| **Lag 1** | 0.87 | 0.79 | 0.65 | 0.24 | 0.04 | 0.19 | 0.35 | 0.62 | 0.58 | 0.46 |
| **Lag 2** | 0.43 | 0.94 | 0.37 | 0.06 | 0.04 | 0.24 | 0.58 | 0.66 | 0.38 | 0.45 |
| **Lag 3** | 0.52 | 0.95 | 0.52 | 0.16 | 0.12 | 0.00 | 0.35 | 0.96 | 0.12 | 0.12 |
| **Lag 4** | 0.75 | 0.90 | 0.19 | 0.31 | 0.30 | 0.01 | 0.37 | 0.95 | 0.21 | 0.03 |
| **Lag 5** | 0.87 | 0.77 | 0.00 | 0.73 | 0.50 | 0.02 | 0.34 | 0.00 | 0.34 | 0.05 |

**Table S7** Granger Causality q values for specific parameter and bacteria

| **Guild** | **Parameter** | **Microbe** | **Lag(p to m)** | **q (p to m)** | **Lag(m to p)** | **q(m to p)** |
|---|---|---|---|---|---|---|
| **methanogen** | Ash Content | Methanofastidiosaceae | 2 | 0.00 | 2 | 0.00 |
| | Ash Content | Methanomassiliicoccaceae | 1 | 0.04 | 2 | 0.00 |
| | Ash Content | Methanosaetaceae | 1 | 0.03 | 2 | 0.00 |
| | ORP | Methanofastidiosaceae | 1 | 0.02 | 4 | 0.04 |
| | ORP | Methanomassiliicoccaceae | 1 | 0.03 | 1 | 0.01 |
| | ORP | Methanosaetaceae | 1 | 0.02 | 4 | 0.03 |
| | pH | Methanofastidiosaceae | 1 | 0.03 | 1 | 0.01 |
| | pH | Methanomassiliicoccaceae | 1 | 0.03 | 1 | 0.00 |
| | pH | Methanosaetaceae | 1 | 0.02 | 2 | 0.03 |
| | VFAs | Methanofastidiosaceae | 1 | 0.03 | 1 | 0.04 |
| | VFAs | Methanomassiliicoccaceae | 2 | 0.02 | 1 | 0.00 |
| | VFAs | Methanosaetaceae | 1 | 0.02 | 5 | 0.04 |
| **acidogen** | VFAs | Lachnospiraceae | 5 | 0.02 | 1 | 0.03 |
| | VFAs | Marinomonadaceae | 2 | 0.05 | 1 | 0.00 |
| | VFAs | Psychromonadaceae | 4 | 0.05 | 1 | 0.00 |
| * p means Parameter, * m means microbial | | | | | | |

### Table S8 Feed-back control policy

| Control Strategy | Trigger (data-driven) | Action (setpoint & dwell) | Purpose (why/ rationale) |
|---|---|---|---|
| Increasing HRT | Any of: EC ≥ 45 mS $cm^{-1}$, ash > 3%, ORP > −125 mV with pH ≤ 5.9 (inhibition window 0–3 d) | Halve outflow for 48–72 h | Avoid premature dilution while short-lag archaeal inhibition completes; divert carbon to VFAs |
| Internal circulation | dVFA/dt ≥ 350 mg COD $L^{-1}$ $d^{-1}$ or ΔMarinomonadaceae ≥ 0.40% (or ≥ 31% $d^{-1}$) or ΔPsychromonadaceae ≥ 0.65% (or ≥ 36% $d^{-1}$) (acidogen window 1–5 d) | 12–18 h boost targeting 30–60 min complete-mix time; then revert | Enhance mass transfer when acidogens lead, converting substrate to VFAs in their window |
| pH banding | Always on (guardbands) | Maintain pH = 5.8–6.0 (floor 5.5, ceiling 6.2); minimal alk feed-forward near draw-offs | Preserve acidogenic regime; prevent souring or drift that erodes yield |
| VFAs removing (10%) | Any of: pH < 5.5; VFA ≥ 12.8 g COD $L^{-1}$ with $\lvert d\text{VFA}/dt \rvert \leq 150$ mg COD $L^{-1}$ $d^{-1}$; pre-emptive: ($d$VFA/$dt$ ≥ 60th pct) and $d^2\text{VFA}/dt^2 < 0$ with rising sentinel taxa | Withdraw 10% VFA-rich liquor; neutral/buffer backfill; cool-down ≥ 24 h before next draw-off | Relieve product inhibition at/near crests; keep production on the rising limb (dominant bottleneck) |

### Table S9 Comparison of VFAs yield between baseline and SW-AAD with feedback control

| Day | VFA_baseline (mg COD/L) | VFA with feedback control (mg COD/L) | ΔVFA (mg COD/L) | Removed VFA (mg COD/L) | :VFA with feedback control (mg COD/L |
|---|---|---|---|---|---|
| 0 | 1037.00 | 1037.00 | 0.00 | 0.00 | 0.00 |
| 1 | 1201.24 | 1201.24 | 0.00 | 0.00 | 164.24 |
| 2 | 1317.00 | 1373.91 | 56.91 | 0.00 | 336.91 |
| 3 | 1392.21 | 1392.21 | 0.00 | 0.00 | 355.21 |
| 4 | 1449.94 | 1449.94 | 0.00 | 0.00 | 412.94 |
| 5 | 1473.00 | 1473.00 | 0.00 | 0.00 | 436.00 |
| 6 | 1377.50 | 1377.50 | 0.00 | 0.00 | 340.50 |
| 7 | 1282.00 | 1282.00 | 0.00 | 0.00 | 245.00 |
| 8 | 2062.73 | 2062.73 | 0.00 | 0.00 | 1025.73 |
| 9 | 3064.00 | 3120.91 | 56.91 | 0.00 | 2083.91 |
| 10 | 3477.55 | 3534.46 | 56.91 | 0.00 | 2497.46 |
| 11 | 3783.50 | 3840.41 | 56.91 | 0.00 | 2803.41 |
| 12 | 3903.00 | 3903.00 | 0.00 | 0.00 | 2866.00 |
| 13 | 3365.50 | 3365.50 | 0.00 | 0.00 | 2328.50 |
| 14 | 2828.00 | 2828.00 | 0.00 | 0.00 | 1791.00 |
| 15 | 6349.22 | 6406.13 | 56.91 | 0.00 | 5369.13 |
| 16 | 9939.00 | 9995.91 | 56.91 | 0.00 | 8958.91 |
| 17 | 10026.96 | 10083.87 | 56.91 | 0.00 | 9046.87 |
| 18 | 10067.06 | 10067.06 | 0.00 | 0.00 | 9030.06 |
| 19 | 10135.00 | 10135.00 | 0.00 | 0.00 | 9098.00 |
| 20 | 10311.06 | 10311.06 | 0.00 | 0.00 | 9274.06 |
| 21 | 10621.00 | 10621.00 | 0.00 | 0.00 | 9584.00 |
| 22 | 11534.21 | 11591.12 | 56.91 | 0.00 | 10554.12 |
| 23 | 12260.00 | 12316.91 | 56.91 | 0.00 | 11279.91 |
| 24 | 11792.81 | 11792.81 | 0.00 | 0.00 | 10755.81 |
| 25 | 10925.19 | 10925.19 | 0.00 | 0.00 | 9888.19 |
| 26 | 10458.00 | 10458.00 | 0.00 | 0.00 | 9421.00 |
| 27 | 11410.27 | 11653.93 | 243.66 | 0.00 | 10616.93 |
| 28 | 12847.00 | 13391.23 | 544.23 | 0.00 | 12354.23 |
| 29 | 13904.23 | 14448.46 | 544.23 | 0.00 | 13411.46 |
| 30 | 14477.00 | 13299.81 | -1177.19 | 1477.76 | 13740.57 |
| 31 | 13946.19 | 13995.01 | 48.82 | 0.00 | 14435.76 |
| 32 | 12870.71 | 12919.53 | 48.82 | 0.00 | 13360.28 |
| 33 | 12026.00 | 12269.66 | 243.66 | 0.00 | 12710.42 |
| 34 | 11564.79 | 12052.11 | 487.32 | 0.00 | 12492.87 |
| 35 | 11339.00 | 11826.32 | 487.32 | 0.00 | 12267.08 |
| 36 | 11723.38 | 11967.04 | 243.66 | 0.00 | 12407.80 |
| 37 | 12473.00 | 12529.91 | 56.91 | 0.00 | 12970.67 |
| 38 | 13682.12 | 13982.69 | 300.57 | 0.00 | 14423.45 |
| 39 | 14526.00 | 15070.23 | 544.23 | 0.00 | 15510.99 |
| 40 | 14048.96 | 14536.28 | 487.32 | 0.00 | 14977.04 |
| 41 | 13163.04 | 13406.70 | 243.66 | 0.00 | 13847.46 |
| 42 | 12686.00 | 12929.66 | 243.66 | 0.00 | 13370.42 |
| 43 | 12816.00 | 13303.32 | 487.32 | 0.00 | 13744.08 |
| 44 | 12946.00 | 12309.29 | -636.71 | 1367.70 | 14117.74 |
| 45 | 12853.96 | 13633.76 | 779.80 | 0.00 | 15442.22 |
| 46 | 12683.04 | 13462.84 | 779.80 | 0.00 | 15271.30 |
| 47 | 12591.00 | 13321.99 | 730.99 | 0.00 | 15130.44 |
| 48 | 12779.00 | 13509.99 | 730.99 | 0.00 | 15318.44 |
| 49 | 12967.00 | 12328.19 | -638.81 | 1369.80 | 15506.44 |
| 50 | 12690.00 | 13226.14 | 536.14 | 0.00 | 16404.39 |
| 51 | 12413.00 | 12949.14 | 536.14 | 0.00 | 16127.39 |

**Table S10 Carbon to VFAS conversion rate.**

| Day | Sample Component | VFAs, mg COD/L | TOC, mg/L | CCE |
|---|---|---|---|---|
| 1 | GLSD Digestate | 1037 | 3820 | 10.18% |
| 7 | | 1282 | 4140 | 11.61% |
| 13 | 50% Digestate + 50% Seaweed | 3903 | 5000 | 29.27% |
| 17 | 25% Digestate + 75% Seaweed | 9939 | 7100 | 52.49% |
| 20 | | 10135 | 7230 | 52.57% |
| 24 | 12.5% Digestate + 87.5% Seaweed | 12260 | 9876 | 46.55% |
| 27 | | 10458 | 11300 | 34.71% |
| 29 | | 12847 | 11300 | 42.63% |
| 31 | | 14477 | 10800 | 50.27% |

**Table S11. Market value of VFAs.**

| VFAs Name | Carbon | Market Value ($/MT) |
|---|---|---|
| Avetic acid | C2 | $650 |
| Propanoic acid | C3 | $2,400 |
| Butyic acid | C4 | $3,300 |
| Pentanoic acid | C5 | $3,500 |
| Hexanoic acid | C6 | $4,100 |
| Heptanoic acid | C7 | $4,600 |

**Table S12. TEA Summary**

| | AD-Biogas | SW-AAD w Control | SW-AAD wo Control |
|---|---|---|---|
| Input feed [ton/day] | 272 | 272 | 272 |
| Retention time [day] | 21.21 | 7 | 7 |
| AD volume [m3] | 5601.09 | 1848.54 | 1848.54 |
| CAPEX digester | 5.39 | 1.99 | 1.99 |
| Capex CHP and gas units | 12.57 | 0.00 | 0.00 |
| Capex VFA units | 0.00 | 5.67 | 5.67 |
| Total capex [million $] | 17.95 | 7.66 | 7.66 |
| OPEX [ utiltiy,labor] | 0.54 | 0.77 | 0.77 |
| CRF | 0.12 | 0.12 | 0.12 |
| Annualized CAPEX | 2.11 | 0.90 | 0.90 |
| Feed cost | - | 3.59 | 3.59 |
| Annual cost | 2.65 | 5.26 | 5.26 |

| | | | |
|---|---|---|---|
| Electricty generation [yearly] | 7128 | 0 | 0 |
| Electrcity reve | 0.57 | 0 | 0 |
| acetic acid [ ton/ yr] | - | 115.90 | 81.73 |
| propanoic acid [ ton/ yr] | - | 41.98 | 29.61 |
| isobutyric acid [ ton/ yr] | - | 14.14 | 9.97 |
| butanoic acid [ ton/ yr] | - | 67.97 | 47.93 |
| isovaleric acid [ ton/ yr] | - | 12.27 | 8.65 |
| pentanoic acid [ ton/ yr] | - | 12.83 | 9.05 |
| Isocrotonic acid [ ton/ yr] | - | 10.63 | 7.50 |
| heptanoic acid [ ton/ yr] | - | 35.34 | 24.93 |
| heptanoic acid [ ton/ yr] | - | 12.65 | 8.92 |
| Rev from VFA sells [ million $] | - | 0.69 | 0.48 |
| Avoided carbon emissions [ ton] | 52.50 | 120.00 | 120.00 |
| Rev carbon credits [ million $] | 0.03 | 0.06 | 0.06 |
| Tipping fee | 4.49 | 3.59 | 3.59 |
| Total revenue | 5.06 | 4.33 | 4.13 |
| Rev without tipping fee | 0.57 | 0.74 | 0.54 |
| Annual Profit | 2.41 | -0.92 | -1.12 |

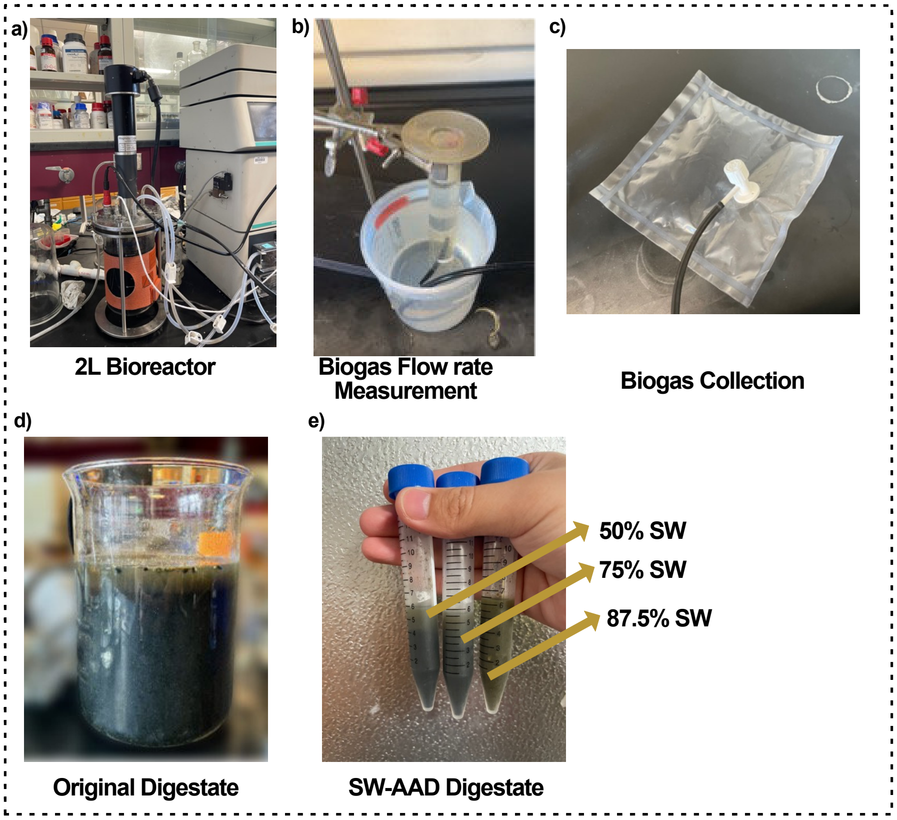


**Figure S1** Photo of a) Bioreactor; b) biogas flow rate measurement; c) biogas collection; d) AD digestate; e) SW-AAD Digestate

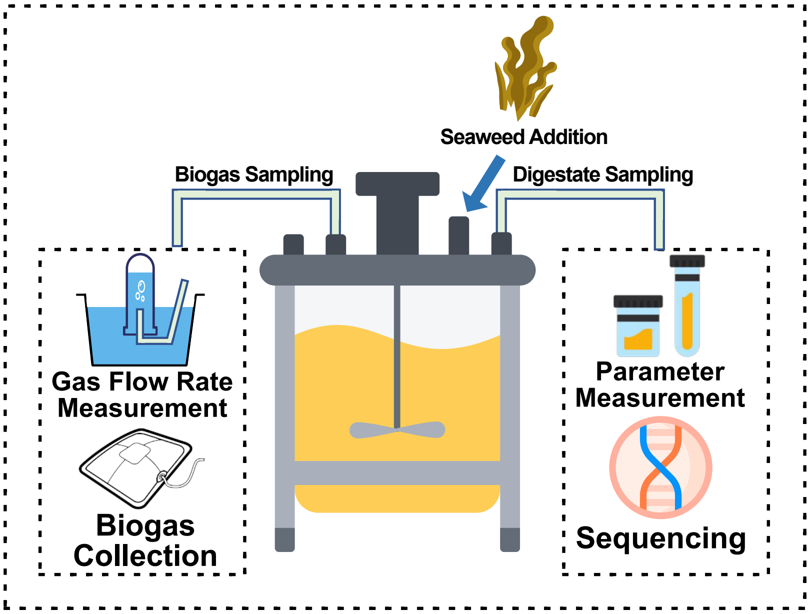


**Figure S2** Diagram of Biogas and Digestate Sampling

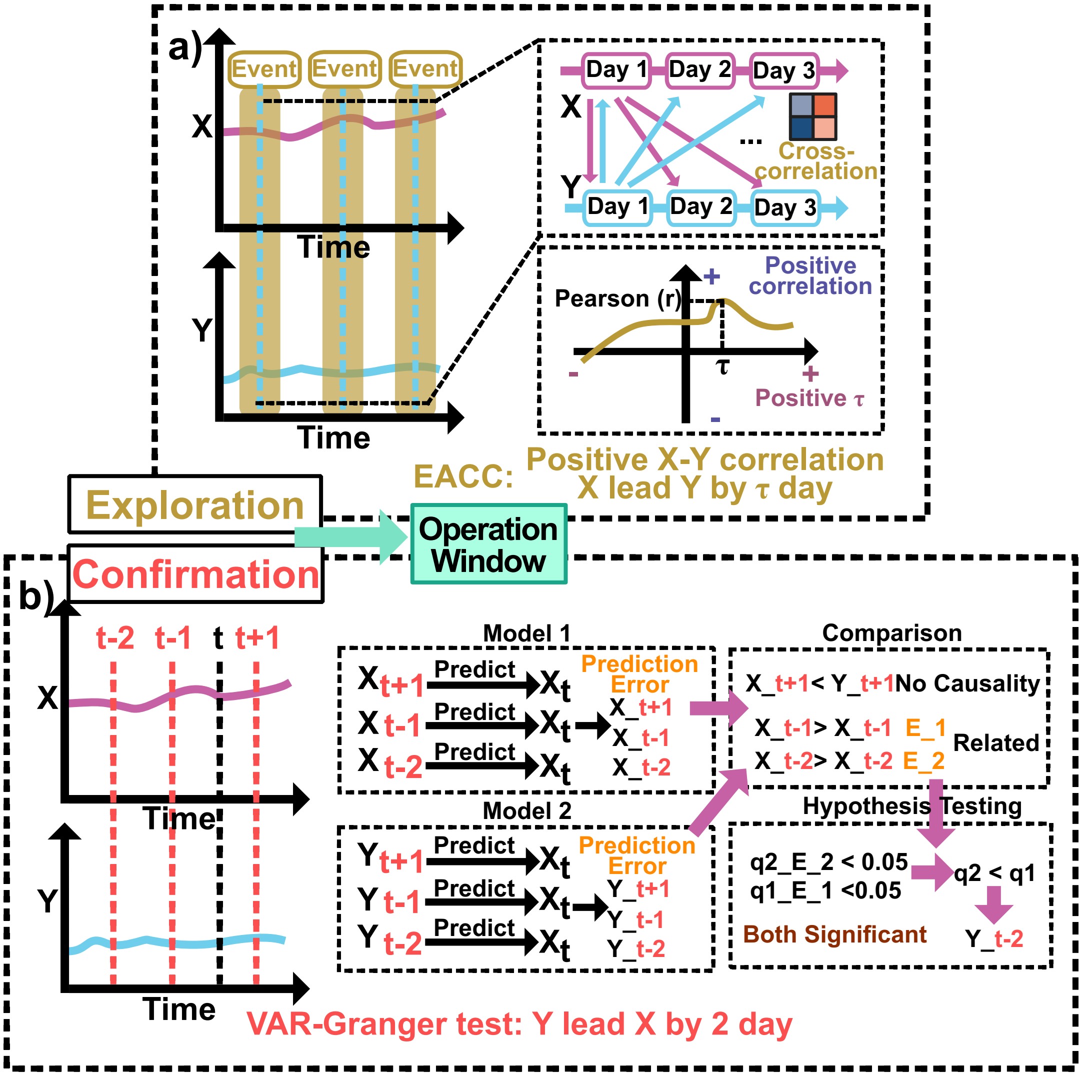


**Figure S3** Methodological Overview of EACC and the VAR-Granger Test. a) Event-Aligned Cross-Correlation (EACC): For each recurring event, a consistent time window is extracted from two time series. The cross-correlation coefficient (r) is calculated for each window across a range of time lags (τ), and these results are then averaged. The final averaged curve reveals the characteristic interplay between the variables: r indicates the strength and direction (positive or negative) of the correlation, while the peak correlation at a specific τ identifies the average time lag of the response. b) VAR-Granger Causality Test: This test determines if the past values of one variable (Y) significantly improve predictions of the future values of another variable (X). It compares two predictive models: Model 1 (Restricted): Predicts X using only the historical data of X. The prediction error is calculated. Model 2 (Unrestricted): Predicts X using the historical data of both X and Y. The prediction error is also calculated.A hypothesis test then determines if the prediction error from Model 2 is significantly lower than that of Model 1. A low q-value (e.g., $q < 0.05$) suggests that Y "Granger-causes" X, meaning its history contains unique information that helps predict X.

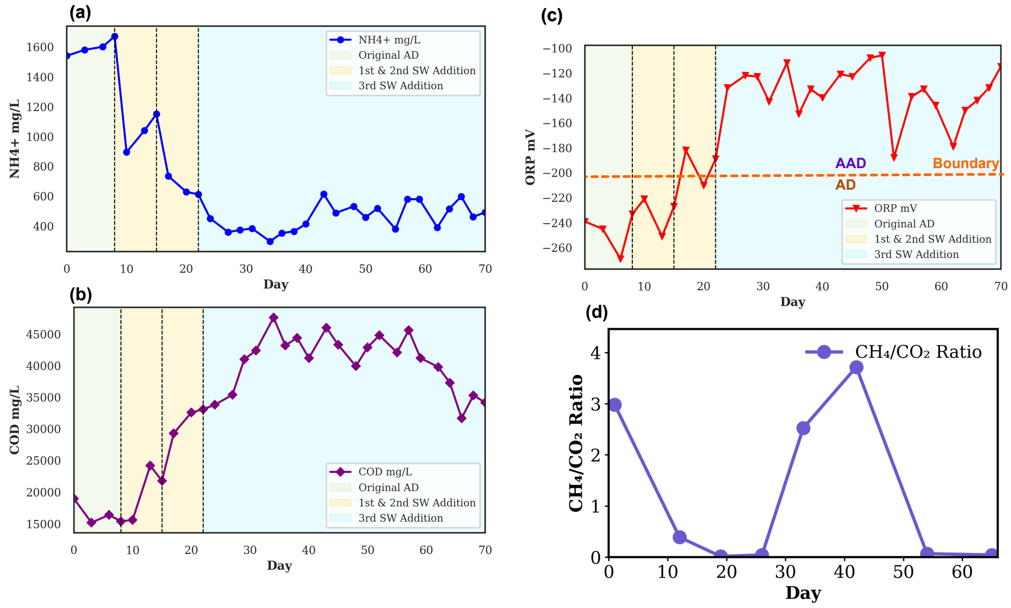


**Figure S4** Operational parameter trend through 3 times brown seaweed replacement with: a) $NH_4^+$ trend; b) COD trend; c) ORP trend and d) $CH_4/CO_2$ Ratio.

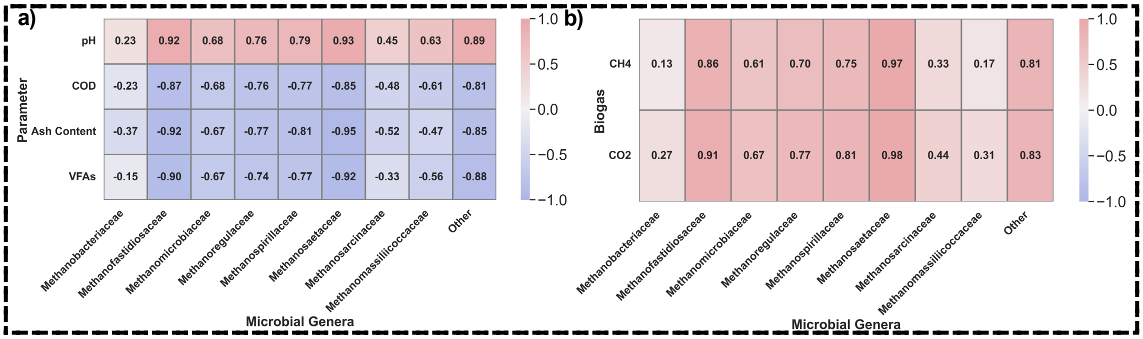

**Figure S5** a) Pearson correlation matrix heatmap (temporal synchrony interplay between engineering parameters and microbial community). VFAs and ash content (salinity proxy) negatively correlate with most methanogens, while pH correlates positively, confirming methanogen suppression under acidic, saline conditions. b) Pearson correlation matrix heatmap (temporal synchrony interplay between methanogenic and biogas). Methanogens exhibit strong positive correlations with $CH_4$ and $CO_2$, affirming their central role in biogas production during the early AD phase and their decline in the SW-AAD phase.

**a)**

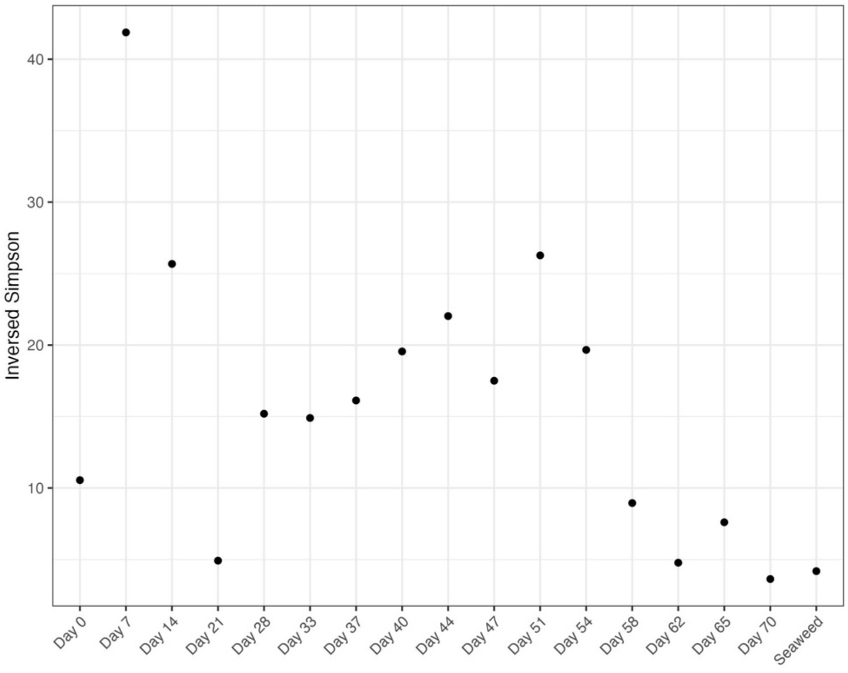


**b)**

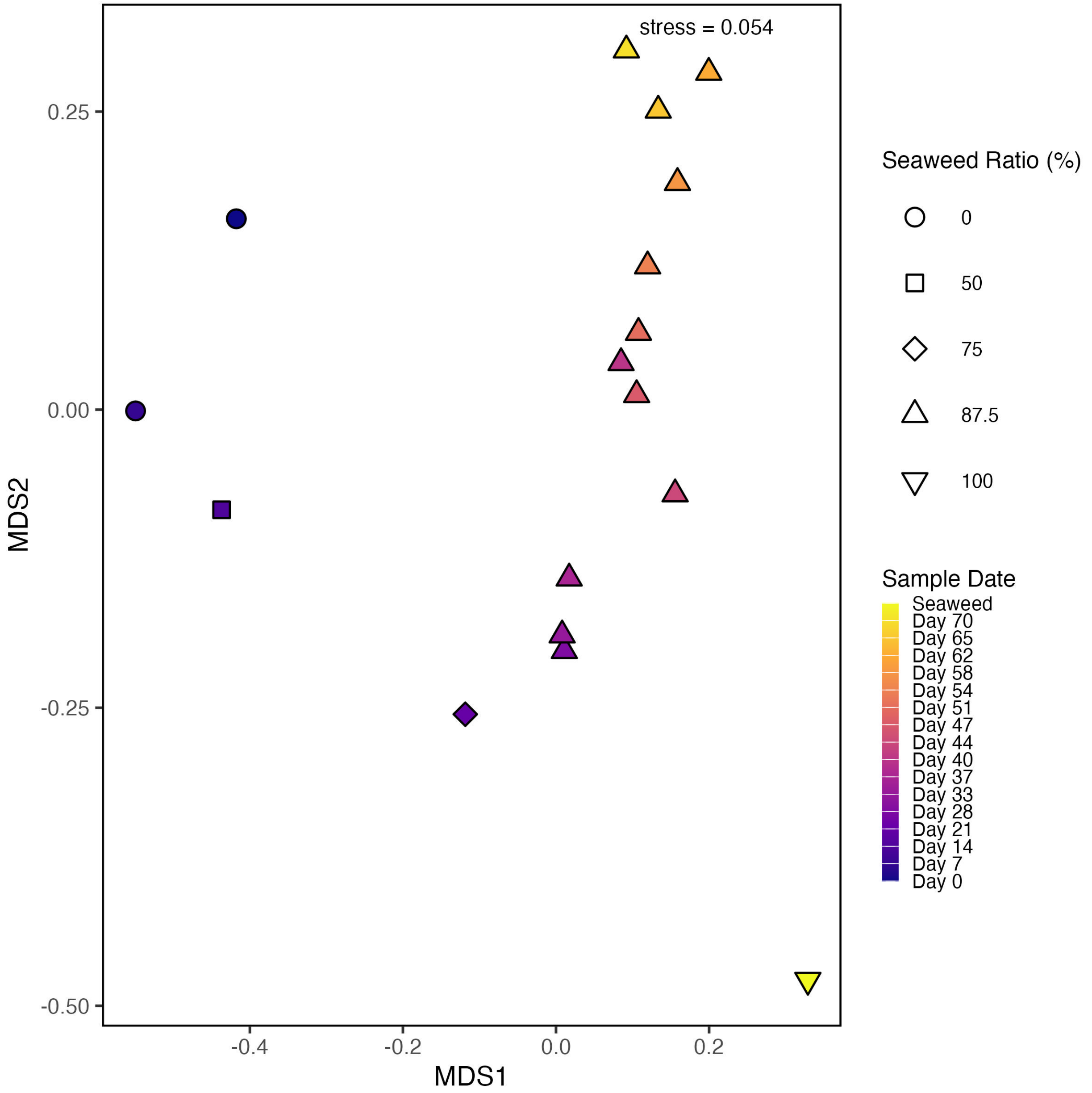


**Figure S6** a. Alpha diversity presenting the inversed Simpson index values throughout the experiment b. Beta diversity was performed with Bray- Curtis dissimilarity profile for OTUs outcomes from 16S rRNA gene sequencing.

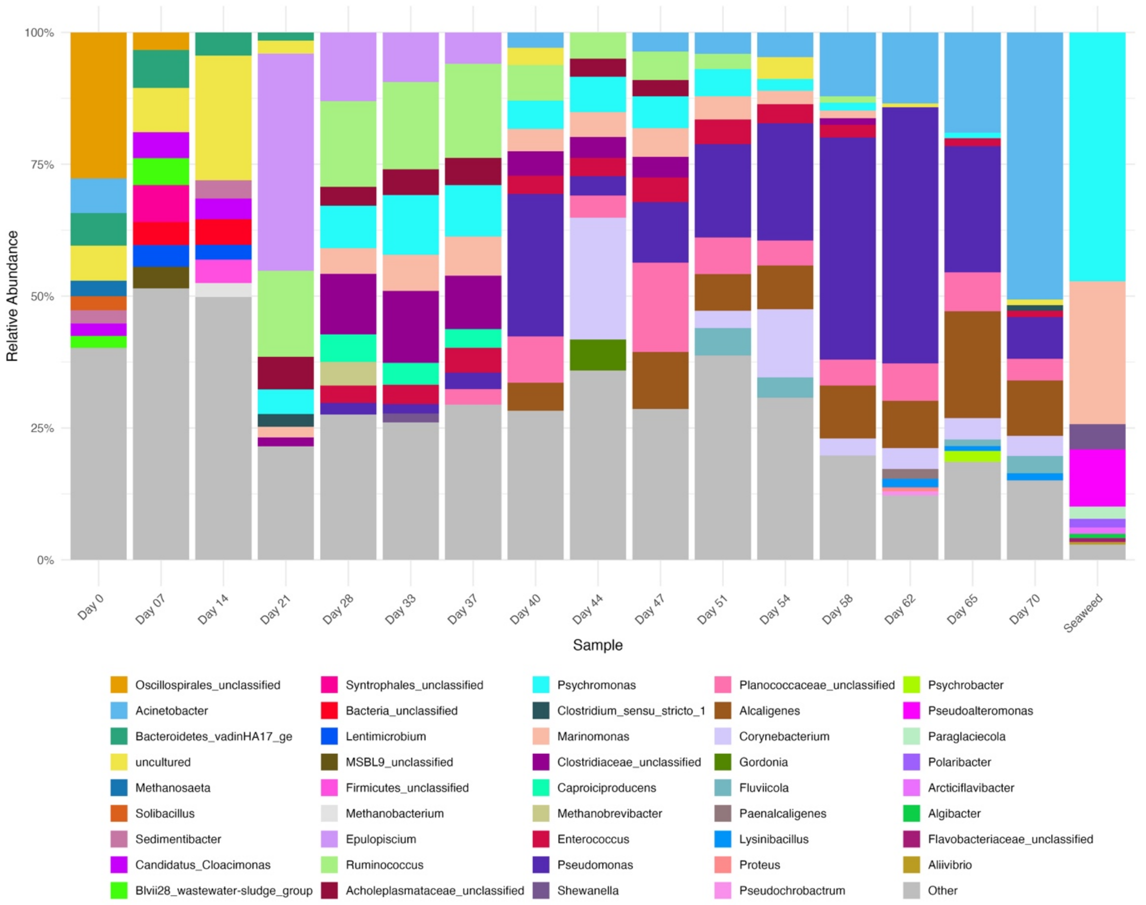


**Figure S7** Top 10 relative abundance of microbiological community at the genus level throughout the trial

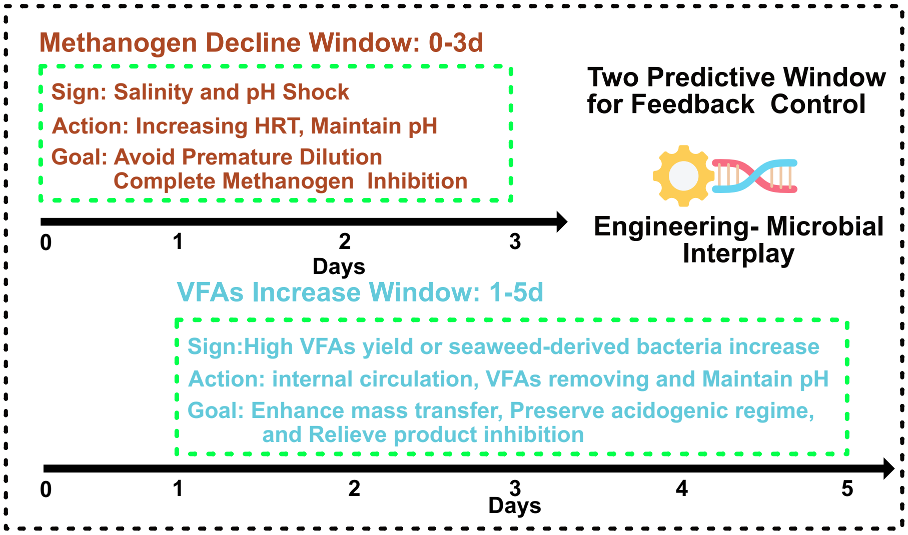


**Figure S8** Two predictive window from asynchrony-resolved feedback control: Mehanogen decline window and VFAs increasing window.

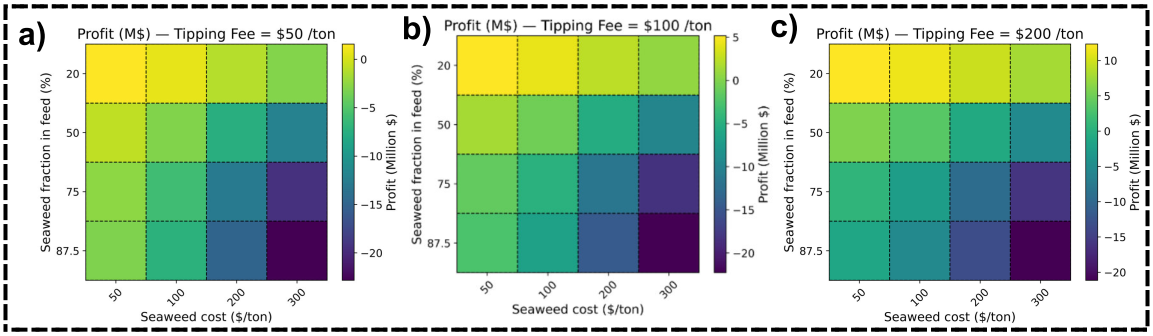


**Figure S9** Sensitivity of process profit (million USD) to seaweed fraction in sludge feed and seaweed cost under different tipping fees of a) 50$/t, b) 100 $/t and b) 200 $/t.

(19) Samanides, C. G.; Koutsokeras, L.; Constantinides, G.; Vyrides, I. Methanogenesis Inhibition in Anaerobic Granular Sludge for the Generation of Volatile Fatty Acids from

CO2 and Zero Valent Iron. *Front Energy Res* **2020**, *8*, 514297. https://doi.org/10.3389/FENRG.2020.00037/BIBTEX.
